# Evaluation of Chromatin Accessibility in Prefrontal Cortex of Schizophrenia Cases and Controls

**DOI:** 10.1101/141986

**Authors:** Julien Bryois, Melanie E Garrett, Lingyun Song, Alexias Safi, Paola Giusti-Rodriguez, Graham D Johnson, Alfonso Buil Demur, John F Fullard, Panos Roussos, Pamela Sklar, Schahram Akbarian, Vahram Haroutunian, Craig A Stockmeier, Gregory A Wray, Kevin P White, Chunyu Liu, Timothy E Reddy, Allison Ashley-Koch, Patrick F Sullivan, Gregory E Crawford

## Abstract

Schizophrenia genome-wide association (GWA) studies have identified over 150 regions of the genome that are associated with disease risk, yet there is little evidence that coding mutations contribute to this disorder. To explore the mechanism of non-coding regulatory elements in schizophrenia, we performed ATAC-seq on adult prefrontal cortex brain samples from 135 individuals with schizophrenia and 137 controls, and identified 118,152 ATAC-seq peaks. These accessible chromatin regions in brain are highly enriched for SNP-heritability for schizophrenia (10.6 fold enrichment, P=2.4×10^−4^, second only to genomic regions conserved in Eutherian mammals) and replicated in an independent dataset (9.0 fold enrichment, P=2.7×10^−4^). This degree of enrichment of schizophrenia heritability was higher than in open chromatin found in 138 different cell and tissue types. Brain open chromatin regions that overlapped highly conserved regions exhibited an even higher degree of heritability enrichment, indicating that conservation can identify functional subsets within regulatory elements active in brain. However, we did not identify chromatin accessibility differences between schizophrenia cases and controls, nor did we find an interaction of chromatin QTLs with case-control status. This indicates that although causal variants map within regulatory elements, mechanisms other than differential chromatin may govern the contribution of regulatory element variation to schizophrenia risk. Our results strongly implicate gene regulatory processes involving open chromatin in the pathogenesis of schizophrenia, and suggest a strategy to understand the hundreds of common variants emerging from large genomic studies of complex brain diseases.

## Introduction

Schizophrenia genomics is progressing rapidly, and our mechanistic understanding of this common and often devastating neuropsychiatric disorder is markedly better than five years ago.^1^ Evidence for a non-specific genetic component for schizophrenia has been known for decades (e.g., via sibling recurrence risk of 8.6 and phenotype heritability estimates of at least 60%).^2–6^ The bulk of the genetic basis of schizophrenia is due to common variation.^7^ A 2014 paper identified 108 genetic regions and a subsequent report has added over 40 new regions, but the implicated regions are broad and usually do not implicate specific genes.^7–9^ It was hypothesized that schizophrenia risk would include many exonic variants of strong effect,^10, 11^ but subsequent large whole exome sequencing studies provide minimal support for this hypothesis.^12–14^ Identifying “actionable” genes has proven complex,^15^ with a few exceptions such as rare exon variants in *SETD1A*^16^ and copy number variation in single genes like *NRXN1* and *C4*.^17–19^

These studies provide strong evidence that genetic risk for schizophrenia results from the concerted effects of many genes. Schizophrenia may be a disorder of subtly altered amounts of protein isoforms rather than changes in individual amino acids. Non-coding regulatory variation is a major contributor to risk for schizophrenia,^20–26^ and genomic regions associated with schizophrenia are enriched for gene expression quantitative trait loci (eQTLs) identified in human brains.^27, 28^ Combining functional genomic data with GWA results may be crucial to deciphering connections to specific genes and disease mechanisms in schizophrenia.^28–30^ ENCODE^31^ and the Roadmap Epigenomics Mapping Consortium^32^ provided considerable human functional genomic data and insights into genomic function. However, these studies provided limited insight into psychiatric disorders as most samples were non-neuronal and none were from affected individuals. This study is part of the PsychENCODE consortium which is intended to provide functional genomic data from the brains of individuals with and without severe neuropsychiatric disorders.^33^

Epigenetic changes in brain are widely hypothesized to partly mediate risk for schizophrenia.^34^ Multiple epigenetic changes have been assessed in schizophrenia (reviewed in^35^), and methylation differences in peripheral blood^36–38^ and brain^39^ have been associated with schizophrenia. Of the many epigenetic changes, chromatin accessibility is particularly important and is a conserved eukaryotic feature characteristic of active regulatory elements, including promoters, enhancers, silencers, insulators, transcription factor binding sites and active histone modifications.^40–43 43–45^ Chromatin accessibility has not been systematically evaluated in human brain for schizophrenia, except for a small study.^46^ Preferentially accessible regions of chromatin can readily be identified by high throughput sequencing following the transposition of sequencing adaptors into the DNA backbone via the Tn5 transposase using the assay for transposase-accessible chromatin sequencing (ATAC-seq).^47^ Unlike other nuclease-sensitivity assays, this approach is amenable to limited amounts of postmortem tissue. As chromatin accessibility can differ by >30% between tissues and cell types,^43^ it is important to study the brains of schizophrenia cases and controls.

Our overall goal is to comprehensively identify active gene regulatory elements in a brain region relevant to schizophrenia, and to quantify how genetic variation alters function such as SNPs that alter chromatin accessibility (i.e., chromatin quantitative trait loci, cQTL). We use these data to parse genetic risk for schizophrenia from large GWA. These analyses provide insight into the molecular mechanisms governing schizophrenia risk. To our knowledge, this is the largest study of chromatin accessibility in schizophrenia and among the largest for any human disease.

## Results

In the ***Online Methods***, we provide the rationale for our choices of ATAC-seq,^47^ brain region, and study design. Key features of our approach include the use of the same samples subjected to mRNA-seq and genotyping analysis by the CommonMind Consortium,^28^ as well as careful experimentation including randomization, blinding, comprehensive quality control, empirical covariate selection, and verification of subject identity.

We performed ATAC-seq on 314 brain samples (142 schizophrenia, 143 control, 23 mood disorders, and 6 other). After quality control, the analysis dataset consisted of ATAC-seq on postmortem Brodmann area 9 (dorso-lateral prefrontal cortex, DLPFC) tissue from 135 cases with schizophrenia and 137 controls (***Figure 1A***). Sixteen individuals with mood disorders were included for open chromatin peak calling and cQTL analyses but otherwise excluded. ***Table 1*** summarizes the demographic and clinical features of the subjects. Cases and controls were comparable for sex, ethnicity, age at death, and postmortem brain pH. Cases had greater postmortem intervals and lower RNA integrity number (RIN) scores relative to controls. These differences appeared to have a lesser impact on DNA-based ATAC-seq assays as cases were comparable to controls for unique aligned reads and normalized open chromatin peak calls. However, these differences motivated comprehensive and careful selection of covariates (***Online Methods***).

**Figure 1:**
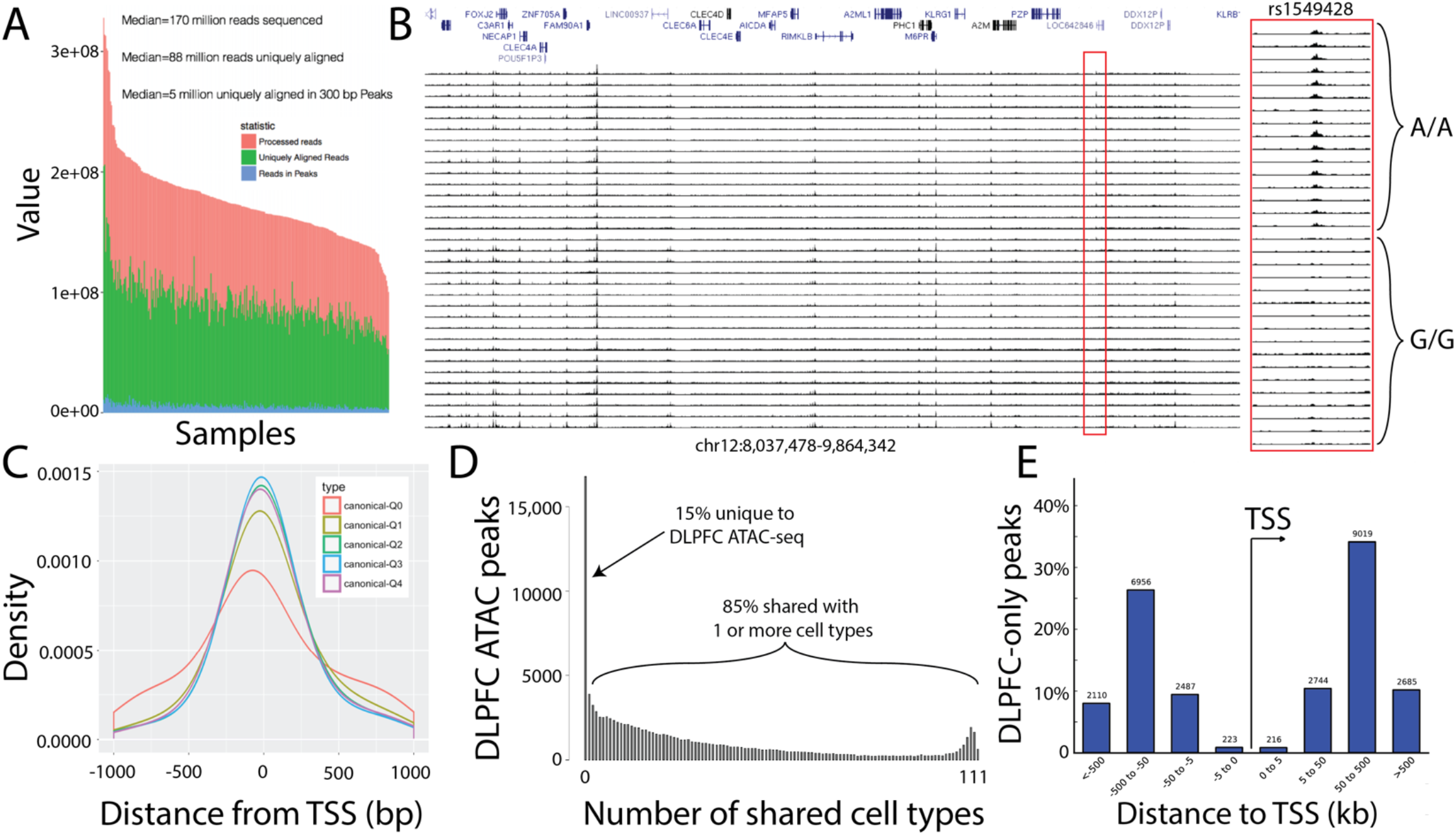
ATAC-seq on frozen DLPFC samples. (A) Sequencing statistics from all libraries show largely similar total number of reads (pink), uniquely aligning reads (green), and reads that map to ATAC-seq peaks (blue). (B) Individual brain ATAC-seq data in a representative genomic region showing largely congruent identification of regions of open chromatin. Some samples have lower signal-to-noise (correlated with covariates like postmortem interval and RNA integrity number). Region in red is a chromatin QTL, note lower signal for indivdiuals with AA genotype vs. GG genotype. (C) Open chromatin in relation to transcription start sites (TSS). The TSS is at zero, right is inward in the direction of transcription, and left is outward from the gene. Density curves for all known GENCODE v25 transcripts (one principal transcript per protein-coding gene). Colored curves show different expression levels in DLPFC: Q0=no expression, Q1-Q4=lowest to highest expression quartiles. (D) Number of DLPFC ATAC-seq peaks that overlap with putative regulatory elements identified from 111 different cell types analyzed by the Epigenome Roadmap Project. Approximately 15% (n=~17,000) ATAC-seq peaks are unique to DLPFC. (E) Location of DLPFC-only ATAC-seq peaks relative to the TSS indicates the majority are non-promoter regions.

**Table 1:** Sample description. All samples were from Brodmann area 9 of left hemisphere. ^†^ Additional ethnicities in cases were African-American (17, 12.6%), Hispanic (5, 3.7%), and Asian (1, 0.74%), and in controls African-American (20,14.6%), Hispanic (16, 11.7%), and Asian (3, 2.2%). ‡ Full details are in reference.^28^ The proportions of oligodendrocytes (0.038), endothelial cells (0.004), microglia (0.0007) were similar in cases and controls. The RNA-based measures are pertinent for the DNA-based ATAC-seq assay as the samples were from the same aliquots.

| Variable | Cases | Controls | Comparison |
| --- | --- | --- | --- |
| Subjects after quality control | 135 | 137 | N/A |
| Male sex, N (%) | 92 (68.2%) | 73 (53.3%) | $\chi^2_1=6.30$, P=0.012 |
| European ethnicity, N (%) † | 112 (83.0%) | 98 (71.5) | $\chi^2_3=7.92$, P=0.048 |
| Age at death, mean (SD) | 73.3 (12.6) | 73.9 (17.7) | $F_{1,271}=0.09$, P=0.76 |
| Postmortem brain pH, mean (SD) | 6.47 (0.24) | 6.50 (0.25) | $F_{1,234}=0.88$, P=0.35 |
| Postmortem brain mass (g), mean (SD) | 1207 (173) | 1155 (166) | $F_{1,269}=6.46$, P=0.012 |
| Estimated cell proportions from RNA-seq ‡ |  |  |  |
| Neurons | 0.781<br>(0.078) | 0.801<br>(0.080) | $F_{1,269}=4.07$, P=0.045 |
| Astrocytes | 0.175<br>(0.067) | 0.156<br>(0.072) | $F_{1,269}=4.39$, P=0.037 |
| Postmortem interval (hours), mean (SD) | 24.3 (15.7) | 10.9 (7.6) | $F_{1,270}=80.2$,<br>P<0.0001 |
| RNA integrity number, mean (SD) | 7.11 (0.79) | 7.57 (0.84) | $F_{1,270}=21.5$,<br>P<0.0001 |
| Unique aligned reads ($\times 10^6$), mean (SD) | 89.12 (17.0) | 89.80 (16.1) | $F_{1,270}=0.12$, P=0.73 |
| Normalized peak calls (FDR 0.01), mean (SD) | 117.3 (70.1) | 130.4 (61.6) | $F_{1,270}=2.66$, P=0.10 |

### ATAC-seq evaluation

ATAC-seq data were aligned and peaks were called using MACS2 (FDR < 0.01; ***Figure 1B***). We identified 118,152 open chromatin peaks totaling 35.5 Mb. As a crude comparison, the human exome has around 131K exons totaling 47 Mb. We compared these regions of open chromatin to those from smaller experiments (***Table S1***). Allowing for small sample sizes, overlap of our open chromatin results with that identified in these other studies was greatest for the most similar studies (DLPFC in adults), somewhat lower in adult cortical samples of sorted neurons, and lower in fetal cortex. Overlap with the diverse ENCODE samples was low, but higher for CNS-relevant samples. These results show congruence of our larger ATAC-seq data with prior experiments, and underscore the need to study brain.

About a quarter of open chromatin regions map near a transcription start site (TSS): 23% were ±5 kb, and 53.0% were >25 kb from any TSS. This distribution was similar to previous studies using DNase-seq.^48, 49^ There is a stronger enrichment for ATAC-seq reads at the TSS for genes that are highly expressed compared to genes that were not expressed (***Figure 1C***). We compared the 118,152 peaks to putative regulatory elements from 111 cell types from reg2map as part of the Epigenome Roadmap project (URLs). We found that 85% of our brain ATAC-seq peaks overlap a promoter or enhancer in one or more of these cell types (***Figure 1D***). This indicates 15% ATAC-seq peak calls are unique to the brain frontal cortex and highlights the need to identify putative regulatory elements from disease relevant tissues. Most of these DLPFC-unique ATAC-seq peaks did not map to promoter regions (***Figure 1E***).

### ATAC-seq peaks are enriched for schizophrenia heritability

We evaluated the relevance of these DLFPC open chromatin regions to schizophrenia using partitioned LD score regression.^50^ This method evaluates whether the SNP-heritability of schizophrenia is enriched in pre-defined genomic features. This approach accounts for multiple technical issues, and conducts a head-to-head comparative evaluation of dozens of genomic features. An earlier analysis found that the SNP-heritability of schizophrenia was strongly enriched in evolutionary conserved genomic regions, but not in open chromatin regions from non-brain cell lines or tissues.^50^ We tested heritability enrichment using the same genomic features^50^ including our DLPFC ATAC-seq peaks and an additional annotation created by expanding the ATAC-seq peaks by ±250 bp to prevent upward bias in heritability estimates.^50, 51^

We found that regions of open chromatin in adult DLPFC were very strongly enriched for genetic variation relevant for schizophrenia (***Figure 2A***): the 1.1% of SNPs located in ATAC-seq peaks explained 11.7% of the SNP-heritability of schizophrenia (10.6 fold enrichment, P=2.4×10^−4^). This level of enrichment was almost as large as conserved regions.^50^ We replicated this result in an independent ATAC-seq dataset from the Chicago psychENCODE group (UIC and U of C) based on 295 adult DLPFC samples from an independent cohort processed with the same bioinformatics pipeline (***Figure 2B*** and ***Figure S1***). We obtained strikingly similar results with 1.5% of the SNPs located in this new set of peaks explaining 13.8% of the SNP-heritability of schizophrenia (9.0 fold enrichment, P=2.7×10^−4^). Thus, common genetic variation that mediates risk for schizophrenia is not randomly distributed in the genome but is concentrated in definable genomic features, particularly regions of open chromatin in brain cortex.

**Figure 2:**
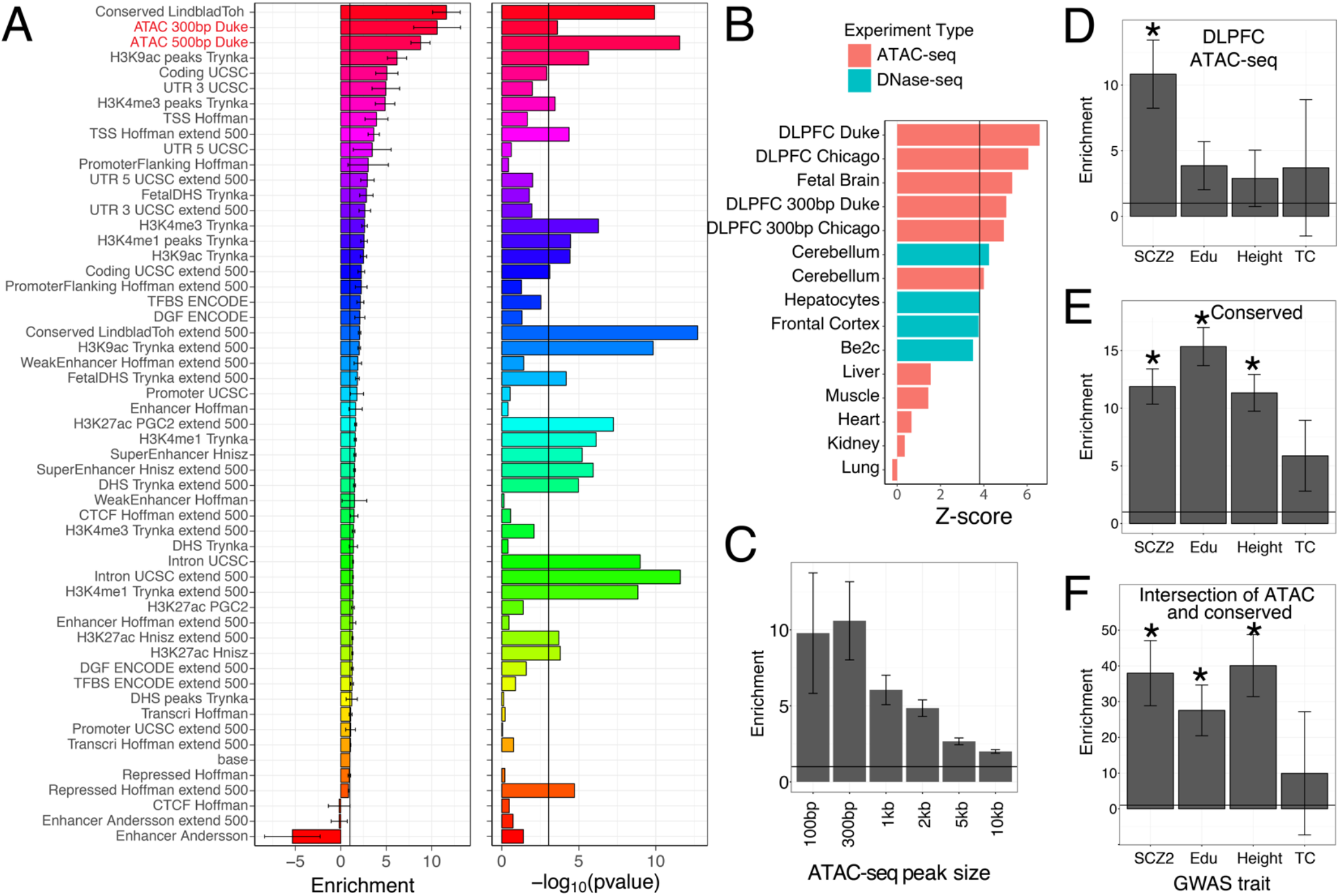
Schizophrenia heritability is enriched for brain-specific accessible chromatin. A) Schizophrenia heritability enrichment (standard error) and significance level (−log10(P)) of different functional genomic and genomic annotations. Enrichment of ATAC-seq regions in DLPFC is second only to genomic regions conserved across 29 Eutherian mammals. B) Enrichment across a subset of 125 DNase-seq and ATAC-seq datasets (see Figure S9 for complete comparison). The black bar represents the Bonferroni significance threshold (=0.01/125). Top enrichments are in DLPFC ATAC-seq peaks generated from Duke and U Chicago groups, followed by other mostly brain specific tissues and cell lines. ATAC-seq tissue samples (cerebellum, liver, muscle, heart, kidney, and lung) represent an independent study to control for batch and method effects C) Schizophrenia heritability enrichment and standard error for peaks of different width. D) Heritability enrichment of ATAC-seq peaks from brain display significant enrichment (* = Bonferroni corrected P < 0.01) for GWA variants associated with schizophrenia, but not for educational attainment (Edu), height, and total cholesterol (TC). E) Heritability enrichment of evolutionary conserved regions are significantly enriched for schizophrenia, educational attainment, and height, but not TC. F) Heritability is 4x more enriched for regions that overlap between ATAC-seq and conservation than for regions that are either conserved or in ATAC-seq peaks.

To evaluate the specificity of this result, we compared open chromatin from DLPFC to that in 124 cell types and tissues generated by the ENCODE Consortium using DNase-seq. The DLPFC ATAC-seq data displayed the greatest association with schizophrenia compared to any other cell or tissue type tested (***Figure 2B*** and ***Figure S2***). These data again indicate the importance of studying relevant tissues from cohort samples. DNase-seq from frontal cortex, cerebellum, a neuroblastoma cell line, and a medulloblastoma cell line also showed more association relative to samples not from brain (***Figure S2***). We generated ATAC-seq from fetal brain samples, and showed slightly lower but still significant enrichment, indicating that relevant regulatory regions that confer risk are accessible earlier in development (***Figure 2B***). Since DNase-seq was used for most of the 124 cell types, we generated ATAC-seq from six diverse human tissues (cerebellum, liver, skeletal muscle, heart, kidney, and lung) and found significant association with schizophrenia only in cerebellum (***Figure 2B***).

The regions flanking ATAC-seq peaks were markedly enriched for schizophrenia heritability. This suggested that causal variants may extend beyond the center of ATAC-seq peaks (***Figure 2A***). This enrichment could be due to our stringent quality control limiting peak size or number, or because open chromatin boundaries vary between individuals.^50^ To test whether SNPs nearer to the peak centers were more likely to be causal, we determined heritability enrichment of ATAC-seq peaks of varying widths (100 bp, 300 bp, 1 kb, 2 kb, 5 kb, and 10 kb). Schizophrenia SNP-heritability enrichment decreased as peak width increased (***Figure 2C***), indicating that SNPs nearer to the peak center explained more SNP-heritability than SNPs farther away.

The SNP-heritability enrichment of DLPFC ATAC-seq peaks was specific to schizophrenia, and was not significantly enriched for GWA variants for educational attainment, height, or total cholesterol (***Figure 2D***). This is in contrast to evolutionarily conserved regions, which are enriched for GWA variants for schizophrenia, educational attainment, and height (***Figure 2E***). To investigate whether sub-regions of the ATAC-seq peaks were particularly enriched for schizophrenia SNP-heritability, we intersected the ATAC-seq peaks with evolutionarily conserved regions,^52^ and found that conserved regions located in DLPFC ATAC-seq peaks were more highly enriched in SNP-heritability (38-fold enrichment; ***Figure 2F***). Conserved regions that map within ATAC-seq peaks cover 6 Mb, much smaller than the 35 Mb covered by ATAC-seq peaks. The intersection of open chromatin and conserved regions tended to be near protein coding transcription start sites (TSS) with 41% located ±10kb from the TSS of the closest protein coding gene whereas only 30.9% of the ATAC-seq peaks and 25.3% of the conserved regions were located ±10kb of a TSS.

Conserved regions in ATAC-seq peaks are significantly enriched for CTCF binding sites (Homer *P*-value < 1 × 10^−100^, MEME-chip evalue < 1 × 10^−100^), which are involved in the formation of topological associated domains,^53^ and other transcription factors involved in neuron differentiation (***Table S2***).^54^ This indicates that common genetic variation implicated in schizophrenia could impact higherorder DNA conformation and/or affect neuron differentiation. Our results support that DLPFC ATAC-seq peaks are enriched for causal variants in schizophrenia, and that restricting these regions to those that are evolutionarily conserved reduces the search space for putative casual variants.

### Identification of differential open chromatin

We next explored possible mechanisms of how these non-coding variants contribute to schizophrenia risk. From an exhaustive list of potential technical and clinical covariates, we identified 13 empirical covariates associated with chromatin peak quantification (***Online Methods, Figure S3***). Controlling for these covariates, we evaluated differential chromatin for age at death and postmortem interval to show that we can detect subtle differences in chromatin accessibility. For age at death, we detected 2,310 sites showing significant differences (FDR < 0.05) in chromatin accessibility (***Figures 3A-B***). These differences may indicate changes in cell heterogeneity (e.g., fewer neurons) or possibly differential chromatin within one or more cell types within the brain that occurs during the aging process. For postmortem interval, we detected 466 sites showing significant differences in chromatin accessibility (***Figures 3C-D***), suggesting that specific regions of the genome are more sensitive to chromatin degradation after death. However, when comparing schizophrenia cases to controls, we detected only three regions with differential open chromatin (***Figures 3E-F***). These three regions did not overlap schizophrenia GWA loci and are not nearby genes that are differentially expressed between cases and controls.^28^ To rule out the possibility of samples with lower signal-to-noise interfering with ability to identify differential chromatin, we performed the same differential chromatin assessment with 62 cases and 75 controls with the highest signal-to-noise, which also resulted in few significant differences in accessibility between schizophrenia and control samples.

**Figure 3:**
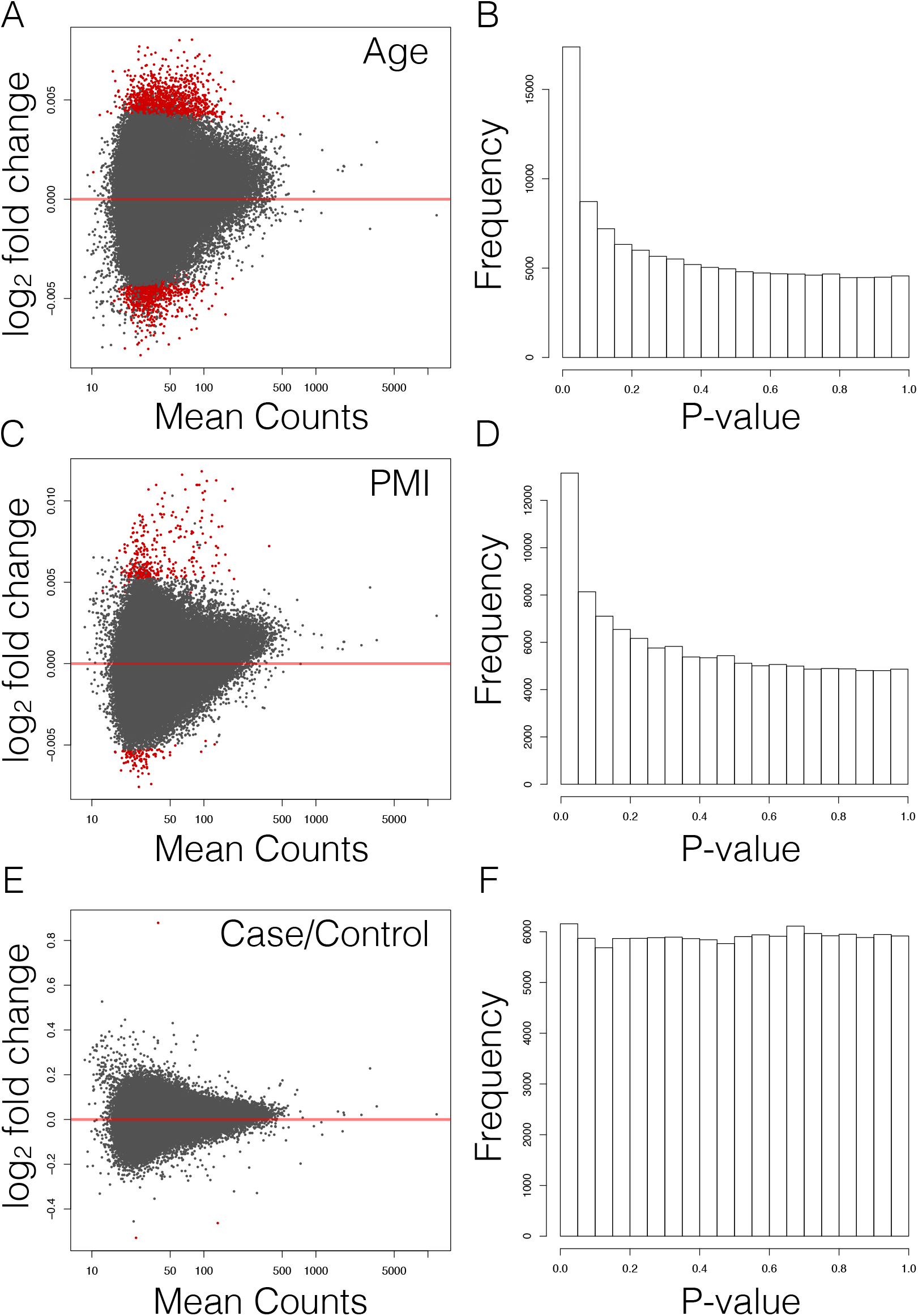
Differential chromatin accessibility differences detected in 288 brain samples. A) Differential chromatin detected as a function of age at death. Dots in red indicate significance (FDR < 0.05). B) Distribution of P-values for age. C) Differential chromatin detected as a function of postmortem interval (PMI). D) Distribution of P-values for PMI. E) Differential chromatin detected as a function of case-control status. F) Distribution of P-values for case-control status. Peaks on chrX were meta-analyzed (inverse variance weighted) and peaks on chrY were only tested in males

A more focused analysis on 1,119 ATAC-seq peaks that map within 96 schizophrenia GWA regions^8^ did not identify any differentially accessible peaks between schizophrenia cases and controls (5% FDR), as well as no enrichment in lower *P*-values. Furthermore, a focused analysis on ATAC-seq peaks located near genes (30 kb upstream to 10 kb downstream) differentially expressed between cases and controls in the same samples^28^ (693 genes, 5% FDR, 4,614 ATAC-seq peaks) did not result in significant differences (5% FDR) and with no enrichment in lower *P*-values. Finally, an analysis of multiple pathways previously associated with schizophrenia, namely genes with RBFOX binding sites, genes whose mRNA bind FMRP, or genes that are the target of antipsychotics^9, 55^, did not result in significant differences between cases and controls (data not shown).

### Identification of chromatin QTL (cQTL)

Previous studies showed that schizophrenia risk alleles are enriched for brain eQTLs.^27^ To determine if genetic variants associated with schizophrenia can also impact chromatin accessibility, we performed a cQTLs analysis using our DLPFC ATAC-seq data (***Figure 4, Online Methods***). In total, we identified 6,200 SNPs that were significantly associated with differences in chromatin accessibility (5% FDR, ***Figure 4A***). There was no skewing of rare or common allele frequency in this subset of SNPs (***Figure 4B***). The influence of gene variation on chromatin accessibility appeared to be localized to nearby regions as 10% of cQTLs (n = 622) were adjacent to (±2 kb) or within the center of the peak with which they were associated (52.0%; ***Figure 4B***).

**Figure 4:**
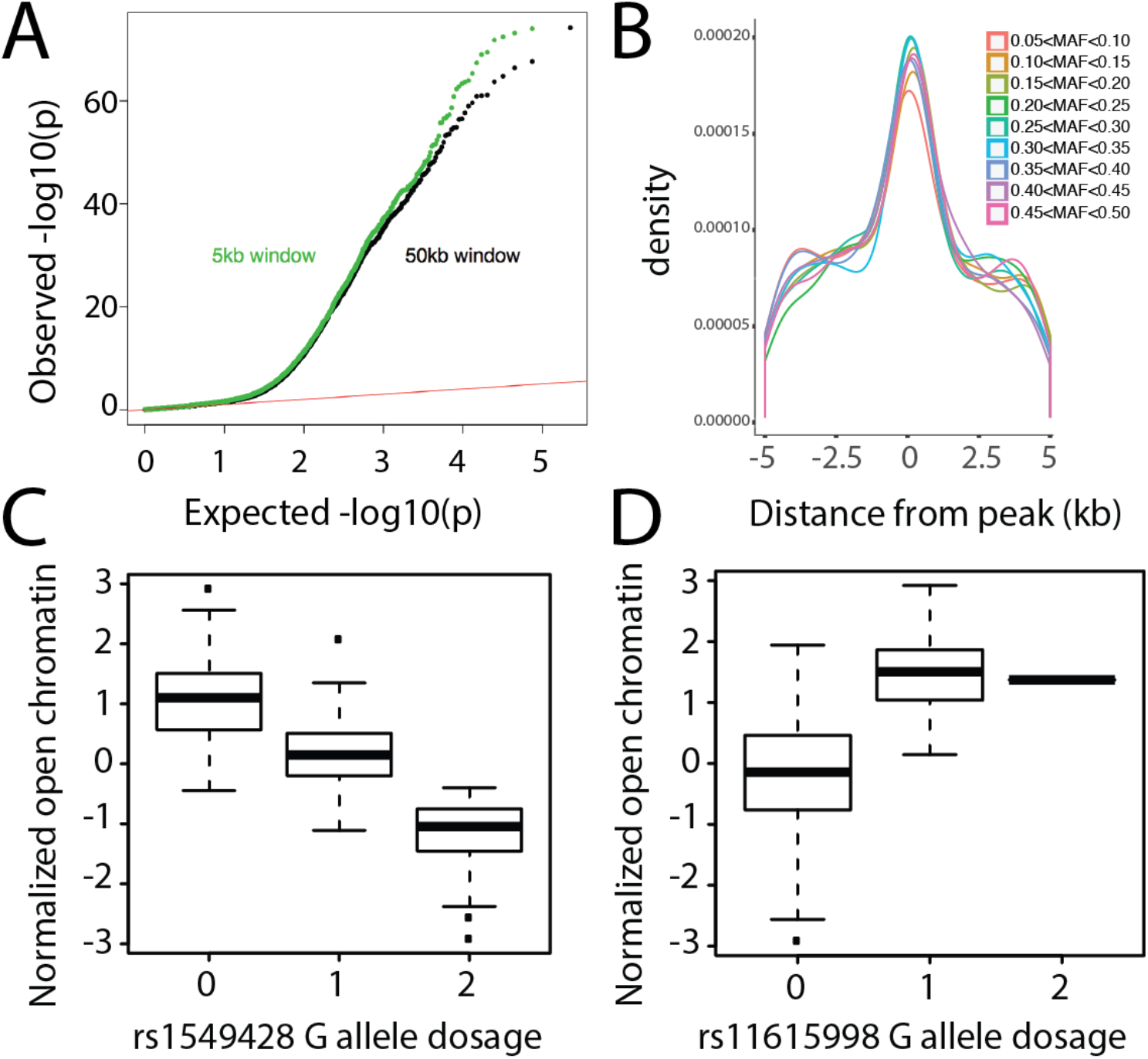
Identification of chromatin QTLs (cQTLs) A) QQ plot of cQTLs using window size of 5kb (green) or 50kb (black). Both show marked deviations from the expected. B) Distance of cQTLs relative to the center of ATAC-seq peaks as a function of minor allele frequency. C) Most significant cQTL, rs1549428 (chr12:9436157–9436457); individuals homozygous for the reference allele (0) display more chromatin accessibility than individuals that are heterozygous (1) and homozygous for the alternate allele (2). D) Most significant cQTL that is also a schizophrenia GWA loci, rs11615998 (chr12:2364960–2365260). Due to a low MAF (0.059) only 1 individual was homozygous for the alternate allele. The results were comparable with or without this homozygous individual.

The most significant cQTL was rs1549428 associated with an open chromatin peak at chr12:9436157-9436457 (*q*=3.9×10^−82^, ***Figure 1B, Figure 4C***). Only 176 of the 6,200 cQTLs (2.8%) were located in a schizophrenia GWA locus,^8^ the most significant of which was rs11615998 in *CACNA1C* (q=2.7×10^−44^, ***Figure 4D***). As an independent confirmation, we show that individuals who are heterozygous for cQTL alleles display allele skewing (***Figure 5A-B***) in the direction that is expected (***Figure 5C***). Similar to eQTLs,^28^ we detect no significant enrichment of cQTLs in schizophrenia GWA regions.

**Figure 5:**
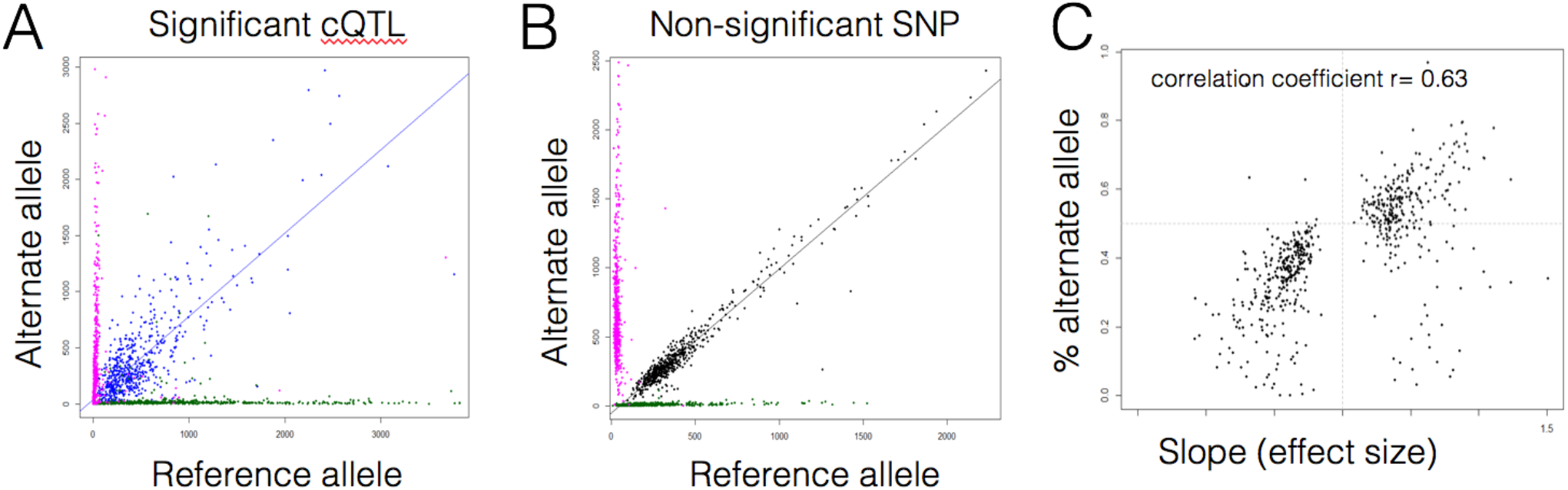
Heterozygous individuals for cQTLs display allele bias. A) Allele counts across cQTLs for individuals homozygous for reference alleles (green), homozygous for alternative alleles (red), or heterozygous for cQTL variants (blue). B) Same as A, but for non-significant cQTLs. C) For individuals heterozygous for cQTLs, we determined the % of each cQTL that correlated to the alternate allele (y-axis) and compared those values to the cQTL effect size (x-axis).

### cQTLs overlap with eQTLs

We compared cQTLs to previously identified eQTLs in the same dataset.^28^ The estimated true proportion of significant cQTLs that are also eQTLs is 23.3% (***Online Methods***), which agrees with similar estimates from lymphoblastoid cell lines.^56^ For SNPs that were both cQTLs and eQTLs, 63.6% were concordant (p < 1 × 10^−4^), meaning the allelic association indicated both more open chromatin and higher gene expression (***Figure 6A***). Despite this correlation of direction of the effect, there was no correlation in the sizes of the effect (***Figure 6B***). Finally, for SNPs in high LD, the direction of effect for cQTL and eQTL was not always the same (***Figure 6C-6D***). Looking at the direction for all QTLs that have pairwise SNPs with r^2^ > 0.8, approximately 31% go in the opposite direction. This is roughly the same percentage going in the opposite direction as it is among all QTLs, regardless of LD (***Figure 6A***). This may indicate independent mechanisms by which these SNPs impact gene expression.

**Figure 6:**
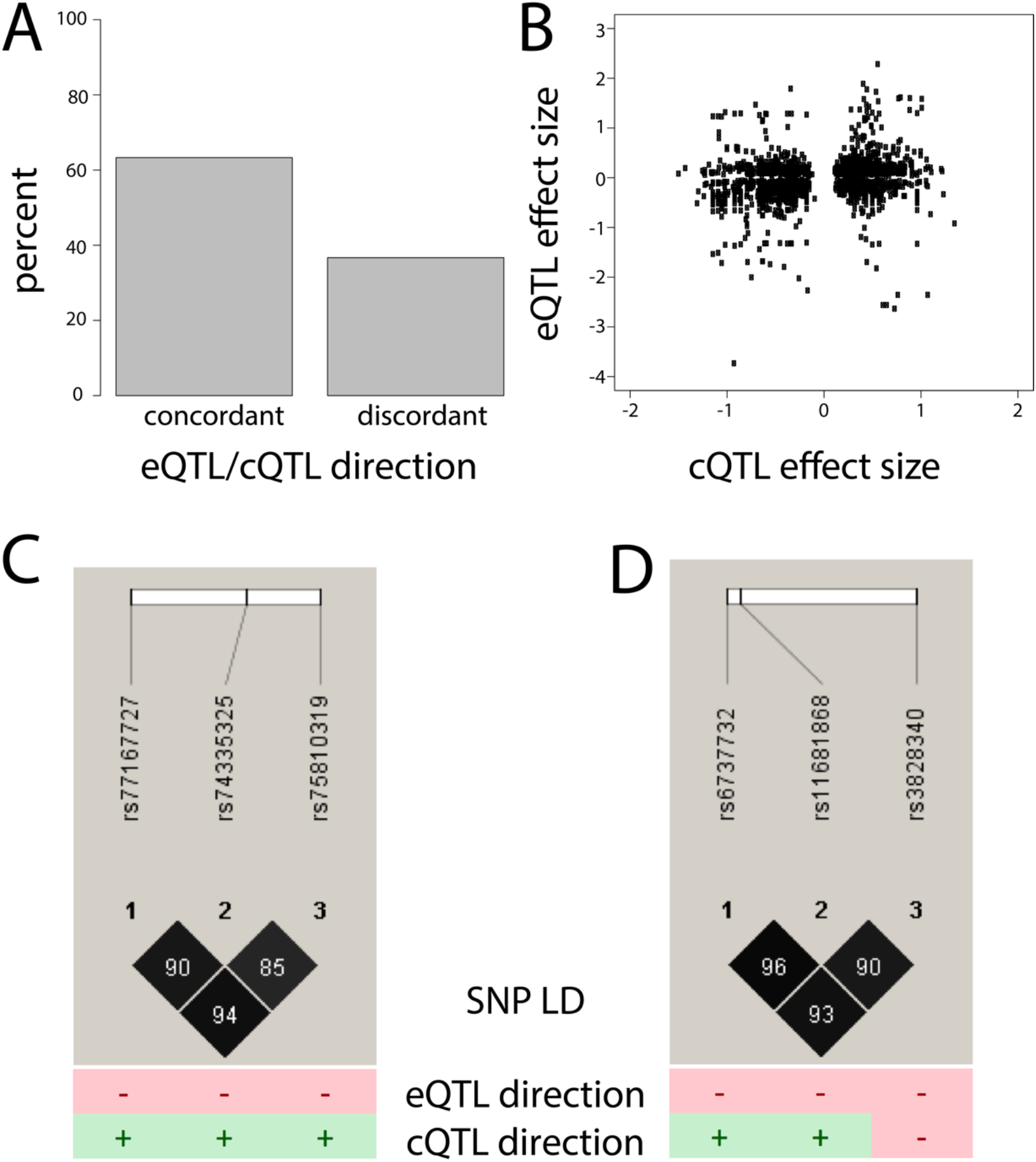
SNPs that are both eQTLs and cQTLs. A) Direction of effects of cQTL vs. eQTL. Concordant effects are when cQTL allele associated with greater open chromatin is also associated with higher eQTL expression. B) Even though the direction of cQTL and eQTL is largely concordant, the effect sizes of these differences are only modestly correlated (r=0.21). C) Three SNPs in high LD on chromosome 1. The tested allele is associated with lower expression and more open chromatin for all three SNPs. Distance between SNP1-2 is 10 kb, and distance between SNP2-3 is 6.3 kb. D) Three SNPs in high LD on chromosome 2. The tested allele is associated with lower expression in all three instances, but with more open chromatin for two SNPs and more closed chromatin for the third SNP. Distance between SNP1-2 is 2kb and distance between SNP2-3 is 20kb.

### cQTL-schizophrenia interaction

Finally, we attempted to identify cQTL SNPs that interacted with schizophrenia status to predict open chromatin regions, but found that none were significant at an FDR below 5%. The number of nominally significant cQTLs in a schizophrenia-associated GWA locus also did not deviate from chance expectations. These results are in line with the eQTL study performed by the CommonMind Consortium that did not detect many eQTLs that interacted with schizophrenia status.^28^

## Discussion

Groups such as the Psychiatric Genomics Consortium have generated some of the largest GWA data for schizophrenia to date.^1^ These studies have largely pointed to non-coding regions of the genome, suggesting that variants in gene regulatory elements contribute to schizophrenia risk. This is supported by whole exome sequencing efforts that have identified few pathogenic coding variants, 14 as well as expression data showing that brain eQTLs are enriched for schizophrenia association statistics.^28^ Thus, we believe that the next step in schizophrenia research is to provide more targeted evidence that schizophrenia risk alleles indeed fall within putative gene regulatory elements, and to characterize the mechanism of those non-coding variants and how they alter gene expression levels. As part of the psychENCODE project,^33^ we have made progress in both areas, and summarize our results in ***Table 2***.

**Table 2:** Summary of results pertaining to schizophrenia genomic findings.

| Empirical findings | Evidence |
| --- | --- |
| Brain open chromatin regions are significantly enriched for schizophrenia SNP-heritability<br>This finding replicated in an independent sample | *Figures 2, S12, S14, S15* |
| Schizophrenia SNP-heritability enrichment in open chromatin is similar to that in regions conserved across 29 Eutherian mammals | *Figure 2A* |
| Regions that are both conserved and open chromatin are particularly enriched | *Figures 2A, 2E, 2F* |
| Many biological features do not show schizophrenia SNP-heritability enrichment (e.g., exons, promoters, other epigenetic marks) | *Figure 2A* |
| Schizophrenia SNP-heritability enrichment is relatively specific and not a feature of GWA generally | *Figure 2D* |
| Adult DLPFC displays similar heritability enrichment relative to fetal PFC | *Figure 2B* |
| Studying adult brain was crucial: higher enrichment compared to 138 tissues/cell types | *Figures 2B, S14* |
| No evidence of open chromatin differences between schizophrenia cases and controls | *Figures 3E-F* |
| Many cQTLs were identified, but no clear relation to schizophrenia | *Figures 4, 5, 6* |

Our study shows that, of diverse genomic and epigenomic datasets that span many different cell types and tissues, ATAC-seq data from DLPFC displays is the most associated with schizophrenia common variant GWA results. Furthermore, genomic regions conserved across 29 Eutherian mammals within adult DLPFC ATAC-seq peaks display an even higher degree of heritability enrichment. This strongly suggests that multiple orthologous data types will be needed to identify causal variants that contribute to schizophrenia risk. This important discovery narrows the search space for identifying causal alleles.

Our initial hypothesis was that these non-coding risk alleles worked through a mechanism that altered chromatin accessibility. This is based on brain eQTLs being enriched for schizophrenia risk alleles,^27^ nearly 600 genes displaying differential expression in brain samples from cases and controls,^28^ and that cQTLs explain a large number of eQTLs in lymphoblastoid cell lines.^56^ Our hypothesis was not confirmed as we did not detect differential chromatin accessibility between schizophrenia cases and controls. This is despite the fact that we were able to detect subtle differences in chromatin accessibility across different age groups and different post mortem intervals, as well as detecting thousands of cQTLs that largely overlap with eQTLs identified in the same samples. Similar to the CommonMind eQTL paper that did not detect many eQTLs that interacted with case/control status,^28^ we did not detect a significant interaction of cQTLs with case/control status. Together, these data indicate that differential gene expression between schizophrenia cases and controls may not be manifested as differential chromatin accessibility.

While it appears that differential chromatin accessibility is not a biological “readout” of non-coding genetic risk for schizophrenia, we cannot rule out that differential chromatin accessibility and/or cQTLs may still be involved. The sample size of our study may have been underpowered to identify differences in chromatin state between cases and controls. We also cannot rule out that cell heterogeneity may mask rare but important cell-type specific signals. In other work we implicated cortical pyramidal neurons as an important cell type for schizophrenia,^57^ and as pyramidal neurons comprise ~40% of the DLPFC, this may be less likely. Furthermore, *in silico* mixing experiments demonstrated that differential chromatin accessibility can accurately be attributed to specific cell types comprising a heterogeneous sample (***Figure S4***). Another possibility is that differential chromatin accessibility between cases and controls is manifested only within specific developmental windows. Indeed, we have identified chromatin accessibility differences that occur during development in post-mitotic mouse neurons.^58^ While careful age-matched comparisons are needed in human to explore this possibility, identifying differences between cases and controls will be challenging as psychotic symptoms typically do not appear until early adulthood.

While these scenarios above are feasible, it is possible that functional non-coding SNPs (including eQTLs) contribute to schizophrenia in a mechanism independent of changes in chromatin accessibility. These functional non-coding variants may still impact specific transcription factor binding sites but overall chromatin accessibility might be maintained and stabilized by other transcription factors and complexes that bind at these regions. Indeed, conserved sequences that are in brain open chromatin regions are highly enriched in CTCF and other transcription factors involved in neuron differentiation. Altering CTCF binding might also impact three dimensional genome structure to allow distal enhancers to abnormally influence gene expression.^59^ Strategies to explore these possibilities include ChIP-seq for specific transcription factors, and Hi-C on brains representing different genotypes. Additional independent strategies include functional detection of variants on regulatory element activity using high-throughput reporter assays like POP-STARR-seq^60^ or regulatory element CRISPR screens.^61^

Future experiments tackling these issues will be needed if we are to further understand the mechanism behind schizophrenia risk as well as disease risk for many other common disorders. These results can help narrow the search space for those studies.

## URLs

BedTools, http://bedtools.readthedocs.io/en/latest/index.html

bowtie2, http://bowtie-bio.sourceforge.net/bowtie2

cutadapt, http://cutadapt.readthedocs.io

MACS2, https://github.com/taoliu/MACS

Mt Sinai NIH Brain and Tissue Repository, http://icahn.mssm.edu/research/labs/neuropathology-and-brain-banking

Open chromatin “blacklists”, https://sites.google.com/site/anshulkundaje/projects/blacklists

Picard tools, https://broadinstitute.github.io/picard

psychENCODE Knowledge Portal, https://www.synapse.org/#!Synapse:syn4921369/wiki/235539

reg2map, https://personal.broadinstitute.org/meuleman/reg2map/HoneyBadger_release

samtools, https://github.com/samtools

UCSC tools, http://hgdownload.cse.ucsc.edu

MEME-chip, http://meme-suite.org/tools/meme-chip

## Acknowledgements

This project was funded by the US NIMH (R01 MH105472, PI Crawford, co-PI Sullivan) and NIGMS (P30GM103328, PI Stockmeier). PFS was supported by the Swedish Research Council (Vetenskapsrådet, award D0886501). JB was supported by the Swiss National Science Foundation. We are deeply indebted to Vahram Haroutunian, Pamela Sklar, and the Common Mind Consortium for providing brain samples. We also thank the NICHD tissue bank (Baltimore, MD) for additional postmortem human tissues, and the NIH NeuroBioBank for fetal brain samples. This work is part of the PsychENCODE Consortium, which was funded by US NIH grants: U01MH103339, U01MH103365, U01MH103392, U01MH103340, U01MH103346, R01MH105472, R01MH094714, R01MH105898, R21MH102791, R21MH105881, R21MH103877, and P50MH106934 awarded to: Schahram Akbarian (Icahn School of Medicine at Mount Sinai), Gregory Crawford (Duke), Stella Dracheva (Icahn School of Medicine at Mount Sinai), Peggy Farnham (USC), Mark Gerstein (Yale), Daniel Geschwind (UCLA), Thomas M. Hyde (LIBD), Andrew Jaffe (LIBD), James A. Knowles (USC), Chunyu Liu (UIC), Dalila Pinto (Icahn School of Medicine at Mount Sinai), Nenad Sestan (Yale), Pamela Sklar (Icahn School of Medicine at Mount Sinai), Matthew State (UCSF), Patrick Sullivan (UNC), Flora Vaccarino (Yale), Sherman Weissman (Yale), Kevin White (U Chicago) and Peter Zandi (JHU). We thank the Duke Sequencing and Genomic Technologies Core facility for sequencing the ATAC-seq libraries.

## Online Methods

### Study design

We conducted a case-control comparison of brain samples to investigate the role of chromatin accessibility in the etiology of schizophrenia. To our knowledge, this is one of the largest studies to date of open chromatin in brain for schizophrenia. Key features were: use of ATAC-seq (a newer method of identifying regions of open chromatin);^47^ use of the same brain tissue samples studied with RNA-seq by the CommonMind Consortium;^28^ and careful experimental design (including randomization, blinding, comprehensive quality control, empirical selection of salient covariates, and verification of subject identity). We attempted ATAC-seq on 314 brain samples (142 schizophrenia, 143 control, 16 bipolar disorder, 7 affective disorder, and 6 other). After the quality control procedures described below, the analysis dataset consisted of ATAC-seq on postmortem brain samples from 135 cases with schizophrenia and 137 controls. For some analyses (e.g., identification of brain cQTLs, covariate selection, differential chromatin analysis), we included 16 individuals with mood disorders. The purpose of including these individuals was to increase power for the cQTLs detection, to increase power to detect covariates affecting ATAC-seq peaks quantification and to better estimate parameters that are not dependent on the case/control status in the differential chromatin analysis.

### Rationale: choice of intact brain versus cell populations

The brain is a complex mixture of cell types.^62^ At present, there is no ideal approach to comprehensively deal with cell heterogeneity. (1) Nuclei sorting from frozen brain provides enrichment for specific cell types.^63–67^ However, sorting is never perfect, relatively few cell types can be sorted in humans, and cells may change state during the 5+ hour process of thawing, dissociation, ultra-centrifugation, and sorting. Sorting neuronal populations from mice with a cell-type specific fluorescent tag is possible, but there is only a ~50% overlap in regulatory regions between mice and human^68^ and Crawford, unpublished). (2) Laser capture microdissection on frozen tissue provides spatial resolution, but yields limited quantity and quality of chromatin, and artifacts from thawing and excess heat are concerns. (3) Single-cell analysis of open chromatin is being developed but is technically difficult, resolution is currently limited, and is not yet available for frozen brain samples.^69^ (4) *Ex vivo* stem cell cultures can yield a realistic cell type but all current *ex vivo* microenvironments do not recapitulate the normal development of the human brain. A further challenge is that the neural cell types that contribute to schizophrenia are not well characterized.^70^

We performed *in silico* mixing experiments and demonstrated that we can detect cell type specific gene regulatory elements at various cell concentrations (***Figure S4A***). Our analysis of the pros and cons of using intact tissues versus cell sorting is shown in ***Figure S4B***. As the cell types causally related to schizophrenia are unknown, we believe that generating chromatin maps in intact brain–i.e., the union of all cell types present in a brain region associated with schizophrenia–provides one strategy of identifying schizophrenia-relevant regulatory elements. Using intact tissue will allow us to analyze more samples, providing better power to identify chromatin QTLs.

### Rationale: choice of brain region

Dorsolateral prefrontal cortex (DLPFC) samples corresponding to, Brodmann areas 9 and 46 were studied. These regions were chosen due to their relevance to schizophrenia based on brain anatomy, imaging, and gene expression.^71–94^ DLPFC was also the focus of a recent paper from the CommonMind Consortium.^28^ In fact, RNA-seq from that study and ATAC-seq data from this study have been generated on tissue aliquots isolated from the same DLPFC dissections.

### Subjects and brain samples

We used ATAC-seq to characterize human DLPFC cortical samples from the Mt Sinai NIH Brain and Tissue Repository (see URLs, Dr. Vahram Haroutunian). All cases met DSM-IV criteria for schizophrenia via standard diagnostic procedures.^28^ Controls had never met criteria for schizophrenia or a psychotic disorder. Subjects were excluded if they had neuropathology related to Alzheimer’s or Parkinson’s disease, acute perimortem neurological insults, or were on a mechanical ventilator near the time of death. No case or control had a large pathogenic copy number variant, and cases had inherited a significantly greater number of schizophrenia risk alleles.^95^

All schizophrenia cases and all controls were dissected from the left hemisphere of fresh frozen coronal slabs cut at autopsy from the DLPFC corresponding to Brodmann area 9). All bipolar disorder samples were from Brodmann area 46. Immediately after dissection, samples were cooled to −190 °C and dry homogenized to a fine powder using a liquid nitrogen-cooled mortar and pestle. Aliquots from each sample were prepared, and used for multiple purposes including ATAC-seq (reported here), RNA-seq,^28^ and SNP genotyping (Illumina OmniExpressExome array). Tissue aliquots were shipped as a dry powder on dry ice to the Crawford lab at Duke University.

Additional adult dorsolateral prefrontal cortex samples (Brodmann area 9, see ***Table S1***) were dissected from postmortem samples from nine adult schizophrenia cases and nine adult control brains, and were obtained from Dr Craig Stockmeier (University of Mississippi Medical Center). Cases and controls were sex- and age-matched. All adult samples were of European ancestry. Controls had no history of psychiatric disorders or substance abuse.

Frontal cortex from nine fetal brains (***Table S1***), gestation age 17-19 weeks, were obtained from the NIH NeuroBiobank (https://neurobiobank.nih.gov). All fetal samples were of African American ancestry. Samples were genotyped on the Illumina Human Omni Express chip in order to confirm sample integrity. Samples were dry homogenized to a fine powder using a liquid nitrogen-cooled mortar and pestle. Aliquots from each sample were prepared, and used for multiple purposes including ATAC-seq, RNA-seq, and DNA microarray. Sample processing was conducted blind to case-control status.

### Confirmation of sample identity

We confirmed subject identity by comparing Illumina SNP genotypes from the Illumina OmniExpressExome array to those recoverable from the ATAC-seq reads (described below). ATAC-seq reads were aligned to hg19 using bowtie2 and variants were called using the multi-sample HaplotypeCaller according to best practices in the Genome Analysis Toolkit.^96^ To achieve a high level of confidence in the variant calls from ATAC-seq, we only kept variants in peaks with a mean read depth ?10 and with minor allele frequency > 0.05. This stringent filtering yielded 10,939 SNPs present in both the ATAC-seq and Illumina data. Identity-by-decent was then estimated for each pairwise combination of ATAC-seq and GWA samples using PLINK^97^ For two subjects, genotypes from ATAC-seq and Illumina genotyping did not match and were excluded in all further analyses.

### ATAC-seq library preparation and sequencing

Samples were processed in batches of eight. Samples were randomly assigned to batches that were balanced with respect to case/control status and sex. Sample processing was conducted blind to case-control status. Frozen pulverized brain samples were received from the Mt. Sinai Brain Repository. Approximately 20 mg of pulverized material was used for ATAC-seq. Frozen samples were thawed in 1 ml of nuclear isolation buffer (20 mM Tris-HCL, 50 mM EDTA, 5mM Spermidine, 0.15 mM Spermine, 0.1% mercaptoethanol, 40% glycerol, pH 7.5), inverted for 5 minutes to mix, and samples were filtered through Miracloth to remove larger pieces of tissue. Samples were centrifuged at 1100 × *g* for 10 min at 4°C. The resulting pellet was washed with 50 μl RSB buffer, centrifuged again, and supernatant was removed. The final crude nuclear pellet was re-suspended in transposition reaction mix and libraries prepared for sequencing as described in Buenrostro et al.^47^ All samples were barcoded, and combined into pools. Each pool contained 8 randomly selected samples (selection balanced by case/control status and sex). Each pool was sequenced on two lanes of an Illumina 2500 or 4000 sequencer (San Diego, CA, USA) at the Duke Sequencing and Genomic Technologies shared resource.

### ATAC-seq initial processing

The raw fastq files were processed through cutadapt (version 1.2.0, URLs)^98^ to remove adaptors and low-quality reads. cutadapt-filtered reads were aligned to hg19 using bowtie2 (version 2.1.0, URLs)^99^ using default parameters. In alignment, all reads are treated as single-read sequences, regardless of whether ATAC-seq libraries were sequenced as single-end or paired-end. The aligned bam files were sorted using samtools (version 0.1.18, URLs),^100^ duplicates removed using Picard MarkDuplicates, and then converted to bed format using BedTools (version: v2.17.0, URLs).^101^ ENCODE “blacklist” regions were removed (i.e., empirically identified genomic regions that produce artefactual high signal in functional genomic experiments, URLs). Narrow open chromatin peaks were called from the final bed files using MACS2, with parameter --nomodel --shift −100 --ext 200. For visualization, bigwig files were generated using wigToBigWig (version 4, URLs)^102^ and bedgraph files were output by MACS2. All data has been submitted and made publicly available on Synapse.

### Identification of sample outliers

We conducted an empirical analysis to identify outliers. An initial analysis identified 8 samples that had only had single end sequencing (unlike the paired end used for all other samples). These samples were excluded.

### Performance of samples

A total of 314 libraries were sequenced across 86 lanes of either Illumina 2500 or 4000, and generated 53,556,161,474 sequences, which total 7,839,829,094,924 bp of data. After filtering with cutadapt, 51,639,643,049 (96.4% of total) sequences were aligned by bowtie2 and generated 27,809,813,130 (53.9% of reads entering aligner) uniquely-aligned reads, and 21,887,340,983 (42.4% of reads entering aligner) multi-aligned reads. Only 3.76% of reads were not aligned. Within the aligned reads, 20,09,897,1743 reads (38.9% of total aligned) were aligned to the mitochondrial genome. In average, MACS2 generated 20,434 ± 12,322 peak calls for each replicate at FDR < 0.01, 28,399 ± 16,917 peak calls at FDR < 0.05 and 35,607 ± 20,750 peak calls at FDR < 0.10. Non-redundant fraction (NRF) of each replicates is 0.881 ± 0.032, PCR Bottleneck Coefficient 1(PBC1) is 0.933±0.030, and PCR Bottleneck Coefficient 2(PBC2) is 19.244 ± 8.295.

### Performance of replicates

We prepared 9 ATAC-seq replicate samples from an independent brain sample aliquots. We observed that the normalized peak values of replicates were significantly more correlated within pairs of replicates (mean spearman correlation = 0.65) then between unrelated samples (mean spearman correlation = 0.42) (P=0.0001, ***Figure S5***).

### Quantification of open chromatin peaks

Peaks called at FDR of 1% in each sample were merged, quantified, and normalized using the diffBind R package.^103^ Only peaks with overlapping coordinates observed in ≥2 samples were quantified. All reads were extended to 300 bp prior to the quantification process. For replicate samples, we retained the replicate with the highest fraction of total reads overlapping with peaks (and for ties, highest number of peaks detected). Peaks were then merged and quantified again as described without the lower quality replicates and forcing the peak width to be 300bp using the summit option in the dba.count function of the diffBind R package. Samples were normalized using the trimmed mean of M values method (TMM).^104^

### Ancestry estimation & population stratification

We wished to capture empirical ancestry using Illumina OmniExpressExome SNP data. These were then available as potential covariates for analyses of differential chromatin accessibility and cQTL. We performed principal component analysis (PCA) of LD-pruned SNP array data using EIGENSOFT.^105^ From the eigenvalue scree plot, we determined that 5 PCs were sufficient to control for effects of population stratification in the GWA data. As such, 5 PCs were included as covariates in the cQTL analysis.

### Evaluation of variables affecting chromatin peaks

For each sample, we recorded a comprehensive set of 206 metadata features that could conceivably capture some aspect of sample quality. These features were collected from the ATAC-seq processing pipeline, RNA-seq processing of aliquots from the same brain regions,^28^ and genome-wide SNP genotyping.^28^ For example, the metadata included transposase batch, date processed, date submitted, PCR cycles, mean GC percentage of sequenced reads, numbers of lanes sequenced, mean mapped read length, subject age at death, sex, diagnosis, postmortem interval, antipsychotic use, history of seizures, and RNA quality. We included 10 ancestry-informative PCs from the genome-wide SNP data. We excluded 19 features with high missingness (e.g., time of death, date of death), 24 features that were invariant in all samples (e.g., ATAC-seq library technician, ATAC-seq data processor, brain region, hemisphere), and 40 features with >5% missing values (e.g., hypertension, body mass index, number of weeks without antipsychotics, tobacco use). To prevent potential over-fitting in downstream analysis, we excluded 18 features with >30 levels (e.g., sequencing batch). We then used the R package mice^106^ to impute missing values using the classification and regression trees methodology. One feature (RNA-seq expression profiling efficiency) could not be confidently imputed and was excluded. This resulted in a total of 104 metadata variables (65 numeric, 39 categorical) for each of the samples. Five variables were deconvolution results estimating the proportions of major cell types in the brain from the RNA-seq data (neuron, astrocyte, oligodendrocyte, microglia, endothelial).^28^

### Covariate selection

In order to detect covariate for our differential chromatin analysis, we performed linear regression of all meta data variables against the first 20 principal components (PCs) of the TMM normalized peak quantification. In an iterative process, we selected one variable (preferentially a variable directly related to the ATAC-seq experiment, explaining one of the largest proportion of variance and with few parameters), regressed its effect on the peak quantifications and performed a new principal component analysis independent of the selected variable(s). We repeated this procedure until we could not further remove the effect of Bonferroni significant variables. Two meta data variables (sequencing lane and whether the sample was excluded from the RNA-seq isolation step) remained associated with PC4, PC9, PC13 and PC15 and could not be corrected because of collinearity with the variable “date submitted”. We believe that not correcting for these two variables is unlikely to result in false positives as they were among the least different meta data variables between cases and controls (96th and 89th among 100 meta variables tested). In total, we selected the following variables for our differential chromatin analysis: GC (%), date submitted, ChrM aligned (%), NRF, RNA-seq intronic aligned (%), RNA-seq astrocyte (%), RNA-seq oligodendrocyte (%), PC1 genotype, seizures, age of death, post-mortem interval, sex, and diagnosis.

### Association between diagnosis and meta data variables

As shown in ***Figure S6***, we observed that many numerical variables were partially correlated within a large cluster of sequencing-related variables (PCR cycles, number of trimmed reads, number of uniquely aligned reads, etc.). To reduce the dimensionality of the data, we applied principal components analysis, and observed that the first principal component largely captured the sequencing-related variables. We also observed that PCs 1, 4, and 9 were significantly associated with diagnosis (***Figure S7***), indicating that the structure of the metadata was different in several dimensions between cases and controls. To identify the metadata variables associated with case-control status, we evaluated the associations of all metadata variables (104) with case-control status (135 cases, 137 controls). We performed linear regression for numerical variables (65) and *χ*^2^ tests for categorical variables (38, excluding diagnosis) using R.

As shown in ***Figure S3***, we observed that 5 numerical and 4 categorical metadata variables were associated at a Bonferroni significance level with case-control status. These variables were: postmortem interval, clinical dementia rating, atypical antipsychotic use, sample storage box, RNA-seq quality, perimortem antipsychotic use, RNA-seq intronic rate, RNA-seq exonic rate, and RNA-seq 28S/16S). The four RNA-seq related variables were highly correlated (***Figure S8***) but not correlated with postmortem interval. The variable most strongly associated with case-control status was postmortem interval (***Figure S9***.

### Association of metadata with ATAC-seq peaks detected

In the analyses above, we did not observe any significant case-control differences in the number of ATAC-seq peaks detected nor in the estimated cell type proportions (from RNA-seq). To identify the variables with an effect on ATAC-seq quality, we performed linear regression for all 100 imputed metadata variables (excluding number of significant ATAC-seq peak calls). We found that 24 variables were associated at a Bonferroni significance level with the number of peaks detected (***Figure S10***). The numerical variables significantly associated with the number of peaks calls formed highly correlated clusters (***Figure S11***), indicating that they captured similar technical variability on the number of peak calls.

Although we did not observe any differences between cases and controls in terms of estimated cell type proportions (from RNA-seq data), we observed that the estimated astrocyte and neuron proportions significantly predicted the number of peak calls. Interestingly, the proportion of neurons and astrocytes were highly correlated with RNA quality (RIN) and slightly less with the mean percentage of GC sequenced. This analysis provided a set of important technical variables affecting the number of peak calls. Nevertheless, this analysis is not sufficient for selecting covariate for the differential chromatin analysis as a variable might not have an effect on the number of peaks detected but an effect on the quantification of the peaks. Hence, we selected covariates for our differential chromatin analysis by looking at the effect of imputed variables on the principal components of the matrix of TMM normalized peak quantification (see above).

### Chromatin QTL analysis (cQTL)

Association analyses for GWA SNPs and open chromatin peaks were evaluated using fastQTL,^107^ which utilizes a ß approximation of permutations to determine significance. Imputed genotype probabilities^28^ were converted to dosages for input into fastQTL. Peaks were normalized and regressed on SNP dosage in a 5kb window, controlling for 10 PCs from PCA of peaks and 5 ancestry PCs from PCA of SNP array data. Only the most significant SNP for each peak was retained. To control for testing multiple peaks, we applied FDR correction^108^ to the ß approximated permutation P-values.

### Heterozygous individuals display allele bias in direction of cQTL association

To provide additional characterization of cQTLs, we analyzed ATAC-seq reads from individuals heterozygous for each of the 600 cQTLs that mapped within an ATAC-seq peak. Allele counts from these individuals show a high degree of allele bias that skews from expected 50:50 split (***Figure 5A***). This is significantly different (Pearson’s Chi-squared test, p-value 2.2e-16) to SNPs that were not identified as cQTLs, which show a higher degree of 50% allele counts (***Figure 5B***). We also find that the direction of the bias in ATAC-seq reads from heterozygous individuals is largely in the same direction as the effect size of each cQTL (***Figure 5C***). In other words, the allele that has more read counts in heterozygous individuals is also the same allele that displays the most accessible chromatin in individuals that are homozygous for that allele. This provides further evidence that cQTL variants are likely contributing directly to chromatin accessibility.

### cQTL interaction with schizophrenia

272 individuals (135 schizophrenia cases and 137 controls) were utilized for the cQTL interaction analysis. This analysis was performed similarly to the cQTL analysis described above, with the addition of a term for the main effect of diagnosis and a term where the SNP genotype interacted with SCZ diagnosis. 10,000 permutations were performed to estimate significance, and the Storey and Tibshirani correction was applied to the exact P-values.^109^

### Overlap of eQTLs with cQTLs

To determine the true proportion of cQTLs that were also eQTLs, we performed a separate FDR calculation using only the SNPs that were present in both analyses, with the qvalue package in R. Because there were multiple eQTLs in a gene, we randomly kept one, matched it to the corresponding cQTL, and performed the FDR correction using that subset of cQTL p-values. We also repeated this procedure twice keeping only the most (round 1) or least (round 2) significant eQTL per gene to obtain a range of estimates for the true proportion cQTLs that were also eQTLs (data not shown). Choosing one random eQTL per gene gave an estimate in between these two extreme estimates, as expected. Conversely, we attempted to estimate the true proportion of eQTLs that were also cQTLs. However, all of the nominal eQTL P-values corresponding to a significant cQTL were < 0.10. Because the distribution was truncated, we could not accurately estimate the true proportion.

### Differential chromatin analysis

We used DESeq2^110^ to detect differential chromatin. The primary analysis was for case-control status, but we also evaluated sex, age at death, and postmortem interval. The following variables were included in the model: GC (%), date submitted, ChrM aligned (%), NRF, RNA-seq Intronic aligned (%), RNA-seq astrocyte (%), RNA-seq oligodendrocyte (%), PC1 genotype, seizures, Age of death, post-mortem interval, sex and diagnosis. All 288 samples were used for this analysis as this increases power to correctly estimate the parameters of the model. The case-control difference was performed between control individuals (N=135) and individuals with schizophrenia (N=137). Although we corrected for sex in all our analysis, we observed that peaks located on the sex chromosomes were more often called significant than peaks located on autosomes. We performed sex stratified analysis for case/control difference in open chromatin and did not observe that peaks on sex chromosomes were enriched in low *P*-values compared to other chromosomes. Therefore, we believe that the initial enrichment in significant hits for all analysis (case/control, sex, PMI and age) on the X and Y chromosomes were likely false positives. In order to prevent potential bias due to the sex chromosomes, we meta-analyzed using an inverse variance weighted approach^111^ for our sex-stratified differential chromatin analysis on chrX and only used *P*-value obtained in male for chrY. For the remaining chromosomes, the full model was used.

### Heritability enrichment analysis

We used LD score regression to estimate heritability enrichment in our ATAC-seq peak calls.^50^ We added SNPs tested in the PGC schizophrenia meta-analysis^8^ and located in our ATAC-seq peak to the “baseline” model which consists of 53 categories representing different genomic annotations (TSS, promoter, enhancer, CTCF binding sites, etc.). In addition, we added SNPs located in a 500bp window (250 bp upstream and 250 downstream) on both side of our ATAC-seq peak calls as an extra annotation to prevent upward bias in the heritability enrichment. Furthermore, we added all SNPs that were present both in the ATAC-seq peak calls and in the conserved regions as an extra annotation. To compare the association of schizophrenia across different tissues and samples, we only added SNPs falling into the respective open chromatin region to the “baseline” model and used the coefficient z-score as a measure of association of the annotation and schizophrenia, as recommended.^50^

### Amount of coverage from and overlap between all DNase-seq/ATAC-seq data

For each DNase-seq and ATAC-seq dataset from 125 tissues, we calculated the total number of bases covered (***Figure S12***) and the average Jaccard index (an indicator of overall similarity of datasets, ***Figure S13***. Jaccard index indicates that ATAC-seq data is most similar to ATAC-seq data from sorted neuronal (NeuN+ sorted) and glial (NeuN-) cells (***Figure S14***).

#### Motif enrichment with MEME-chip

We intersected conserved regions^52^ with our ATAC-seq peaks. As MEME-chip requires all input sequences to have the same length, we set the width of each intersected region to 32bp (16bp upstream and downstream of the center of the intersected region, corresponding to the mean size of the intersected regions). We obtained hg19 sequence corresponding to these regions using bedtools. We then used MEME-chip^112^ (URL) to look for motif enrichment using default settings;; and used HOMER^113^ (v4.9) with intersected conserved regions with our ATACseq peaks as input and randomly chosen 1000,000 conserved regions as background to do motif search.

### Data sharing

These ATAC-seq data are available from Sage Bionetworks-Synapse website via the psychENCODE Knowledge Portal (URLS).

### Genome build

All genome coordinates are GRCh37 / hg19.

---

**Table S1.** Overlap of ATAC-seq data from this study with external samples. CNS=central nervous system. DHS=DNase I hypersensitivity. NeuN is a marker for post-mitotic neurons. This table compares our ATAC-seq dataset with regions of open chromatic identified in other experiments including ENCODE,^31^ Roadmap Project,^32^ a pilot study from Roussos and colleagues,^46^ and a separate pilot study from our team (unpublished). Given the widely varying sample sizes and different methods used, we queried the proportion of the open chromatin overlapping with our study. Overlap of open chromatin identified in these other studies was greatest for the most similar studies (DLPFC in adults), somewhat lower in adult samples of sorted neurons, and considerably lower in fetal cortex. Overlap of ENCODE data was low for the full set of highly diverse tissues and cell lines but somewhat higher for the set of CNS-relevant tissues and cell lines. This pattern of results suggests the congruence of our data with those from these other studies.

| Source | Technique | Tissue | Samples | % Found | Peaks |
| --- | --- | --- | --- | --- | --- |
| ENCODE | DHS | Highly diverse | 124 | 5.48 | 2815428 |
| ENCODE | DHS | CNS-relevant tissue & cell lines | 20 | 13.42 | 642219 |
| MSSM Roussos | ATAC-seq | Brain, PFC, adult, sorted NeuN- | 8 | 27.97 | 115216 |
| MSSM Roussos | ATAC-seq | Brain, PFC, adult, sorted NeuN+ | 8 | 35.97 | 115216 |
| Roadmap Project | ATAC-seq | Brain, frontal, fetal | 2 | 14.36 | 589344 |
| Sullivan-Crawford pilot | ATAC-seq | Brain, frontal, fetal | 9 | 6.27 | 1118135 |
| Sullivan-Crawford pilot | ATAC-seq | Brain, DLPFC, adult, SCZ cases | 9 | 43.71 | 99439 |
| Sullivan-Crawford pilot | ATAC-seq | Brain, DLPFC, adult, controls | 9 | 72.33 | 49969 |

**Table S2.**
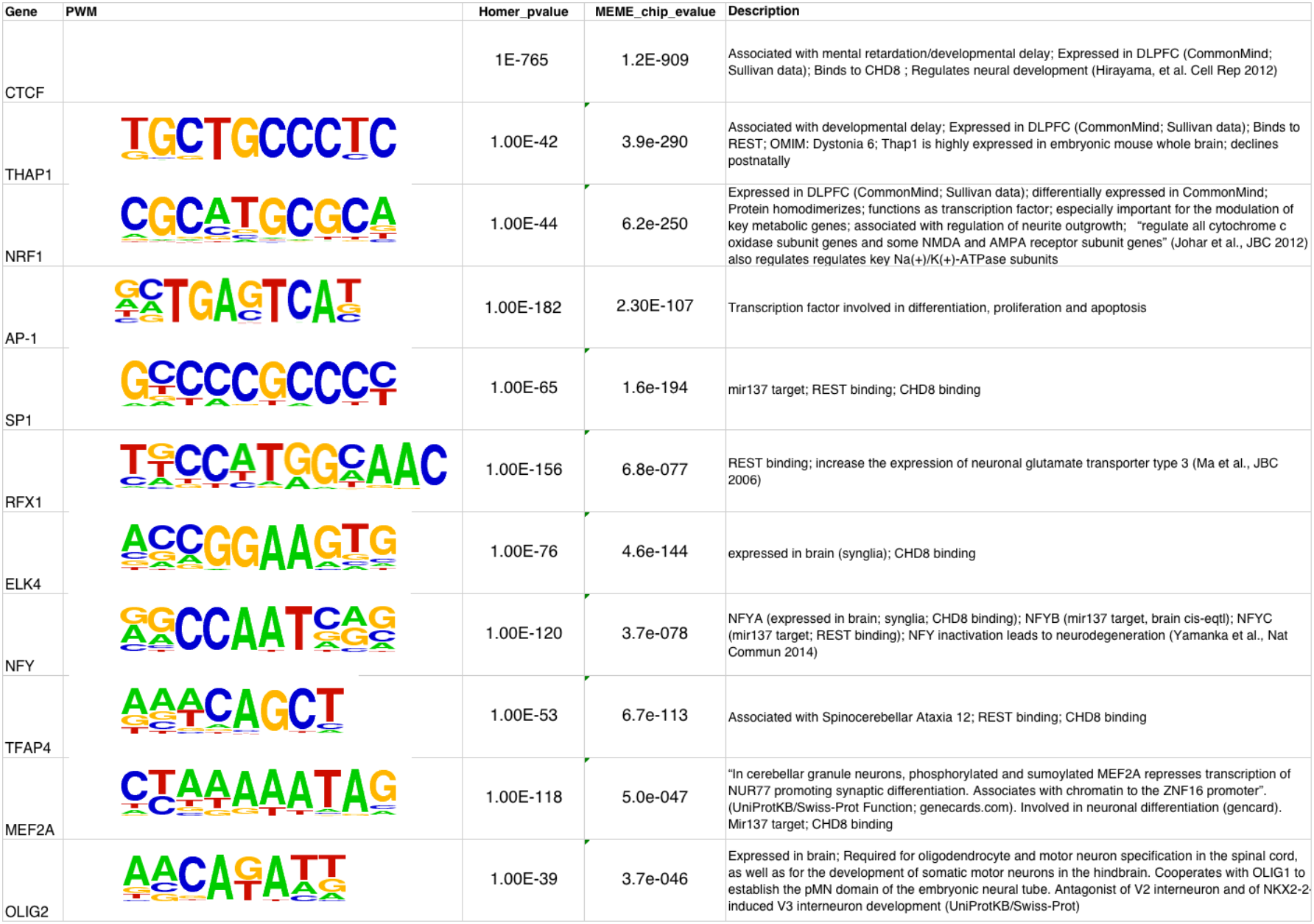
Enrichment of transcription factor binding motifs in DLPFC ATAC-seq peaks that overlap evolutionarily conserved sequences. Shown are top enriched motifs using two different motif enrichment methods, Homer and MEME-chip.

| Gene | PWM | Homer_pvalue | MEME_chip_evalu | Description |
| --- | --- | --- | --- | --- |
| CTCF |  | 1E-765 | 1.2E-909 | Associated with mental retardation/developmental delay; Expressed in DLPFC (CommonMind; Sullivan data); Binds to CHD8 ; Regulates neural development (Hirayama, et al. Cell Rep 2012) |
| THAP1 |  | 1.00E-42 | 3.9e-290 | Associated with developmental delay; Expressed in DLPFC (CommonMind; Sullivan data); Binds to REST; OMIM: Dystonia 6; Thap1 is highly expressed in embryonic mouse whole brain; declines postnatally |
| NRF1 |  | 1.00E-44 | 6.2e-250 | Expressed in DLPFC (CommonMind; Sullivan data); differentially expressed in CommonMind; Protein homodimerizes; functions as transcription factor; especially important for the modulation of key metabolic genes; associated with regulation of neurite outgrowth; "regulate all cytochrome c oxidase subunit genes and some NMDA and AMPA receptor subunit genes" (Johar et al., JBC 2012) also regulates regulates key Na(+)/K(+)-ATPase subunits |
| AP-1 |  | 1.00E-182 | 2.30E-107 | Transcription factor involved in differentiation, proliferation and apoptosis |
| SP1 |  | 1.00E-65 | 1.6e-194 | mir137 target; REST binding; CHD8 binding |
| RFX1 |  | 1.00E-156 | 6.8e-077 | REST binding; increase the expression of neuronal glutamate transporter type 3 (Ma et al., JBC 2006) |
| ELK4 |  | 1.00E-76 | 4.6e-144 | expressed in brain (synglia); CHD8 binding |
| NFY |  | 1.00E-120 | 3.7e-078 | NFYA (expressed in brain; synglia; CHD8 binding); NFYB (mir137 target, brain cis-eqtl); NFYC (mir137 target; REST binding); NFY inactivation leads to neurodegeneration (Yamanka et al., Nat Commun 2014) |
| TFAP4 |  | 1.00E-53 | 6.7e-113 | Associated with Spinocerebellar Ataxia 12; REST binding; CHD8 binding |
| MEF2A |  | 1.00E-118 | 5.0e-047 | "In cerebellar granule neurons, phosphorylated and sumoylated MEF2A represses transcription of NUR77 promoting synaptic differentiation. Associates with chromatin to the ZNF16 promoter". (UniProtKB/Swiss-Prot Function; genecards.com). Involved in neuronal differentiation (gencard). Mir137 target; CHD8 binding |
| OLIG2 |  | 1.00E-39 | 3.7e-046 | Expressed in brain; Required for oligodendrocyte and motor neuron specification in the spinal cord, as well as for the development of somatic motor neurons in the hindbrain. Cooperates with OLIG1 to establish the pMN domain of the embryonic neural tube. Antagonist of V2 interneuron and of NKX2-2; induced V3 interneuron development (UniProtKB/Swiss-Prot) |

## Figures and Supplemental Figures

Bryois et al.

### Supplemental Figures

**Figure S1:**
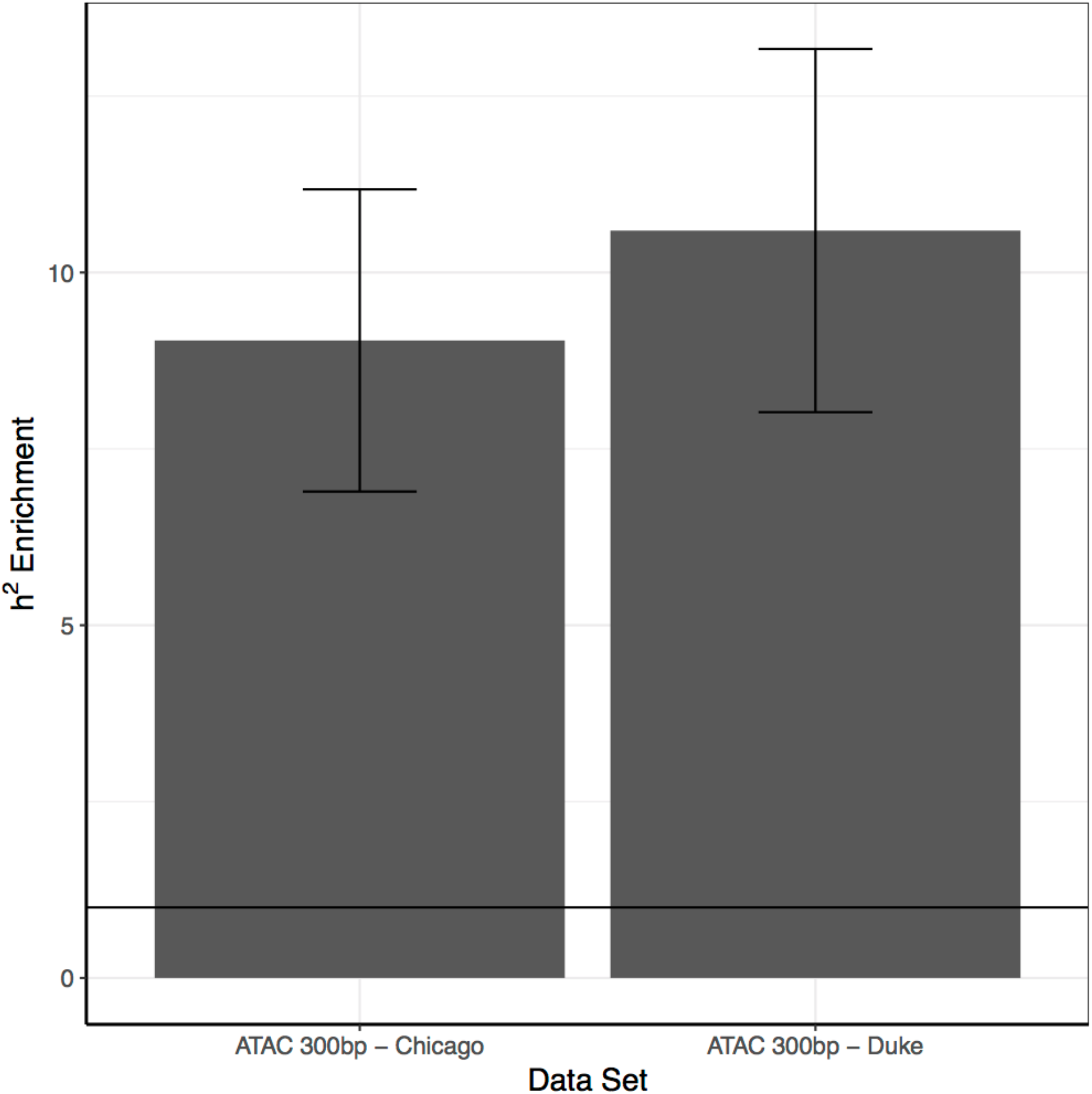
Heritability enrichment for schizophrenia. For the 118,152 ATAC-seq peaks analyzed in this study (Duke University) and replication in an independent ATAC-seq data set consisting of 157,660 peaks obtained using the same bioinformatics pipeline (University of Chicago). The heritability enrichments were obtained using partitioned LD score regression using the full baseline model, the ATAC-seq peaks and an additional track with the ATAC-seq peaks extended on both sides by 250bp. Only the heritability of the ATAC-seq peaks is shown here.

**Figure S2:**
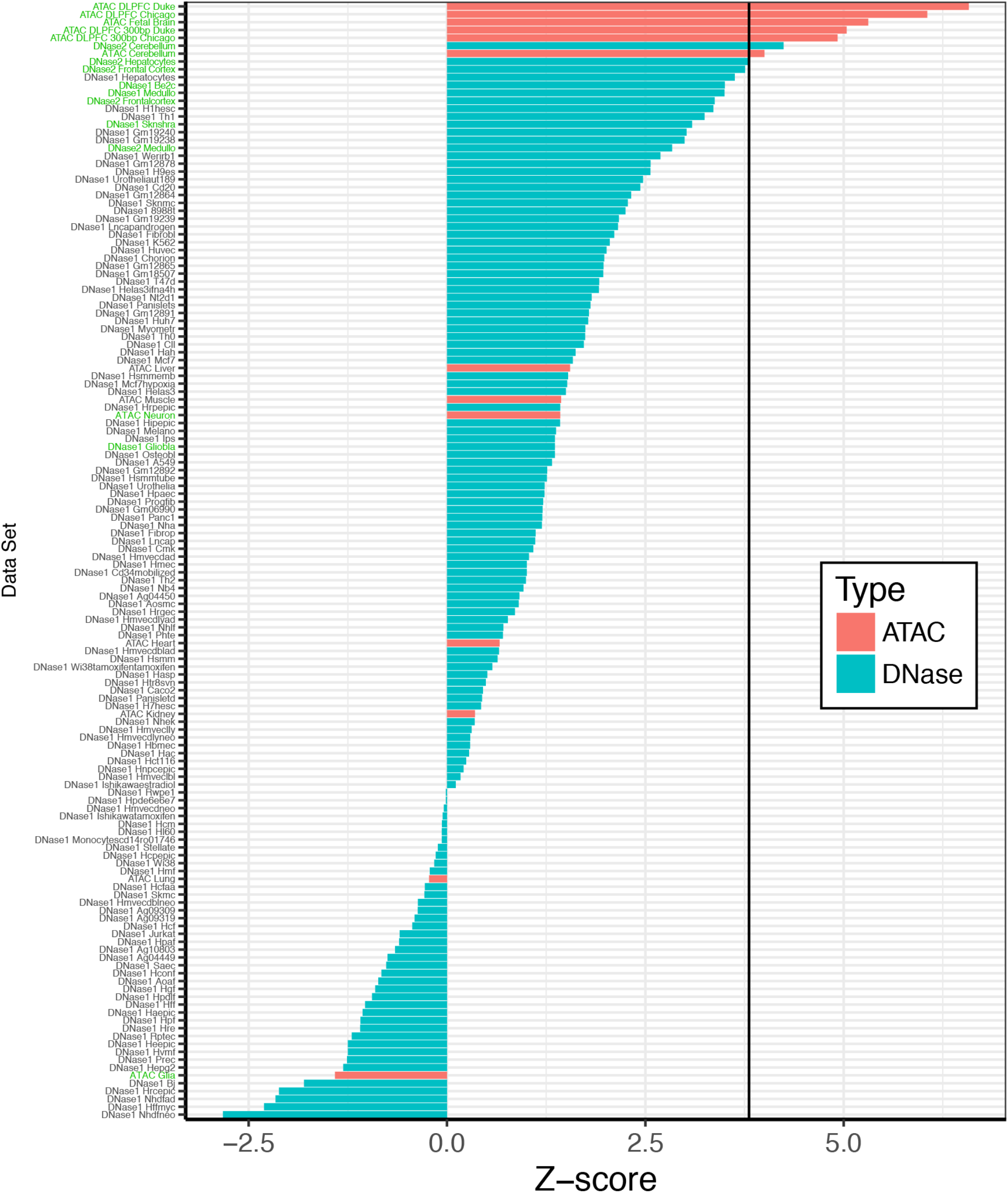
Heritability enrichment from DNAse-seq and ATAC-seq across 142 cell and tissue types. Heritability enrichment Z-scores were determined from ATAC-seq (red) DNase-seq (blue) data generated from 142 cell types and tissues, including ATAC-seq data generated by Duke and Chicago (University of Illinois at Chicago, and the University of Chicago) using partitioned LD score regression. Each dataset was added to the full baseline model and tested one at a time. The black bar represents the Bonferroni significance threshold (p adjusted =0.01). Note that the top enrichment values were from ATAC-seq samples from Duke and University of Chicago. Cell lines and tissues that were from brain origin are listed in green text.

**Figure S3:**
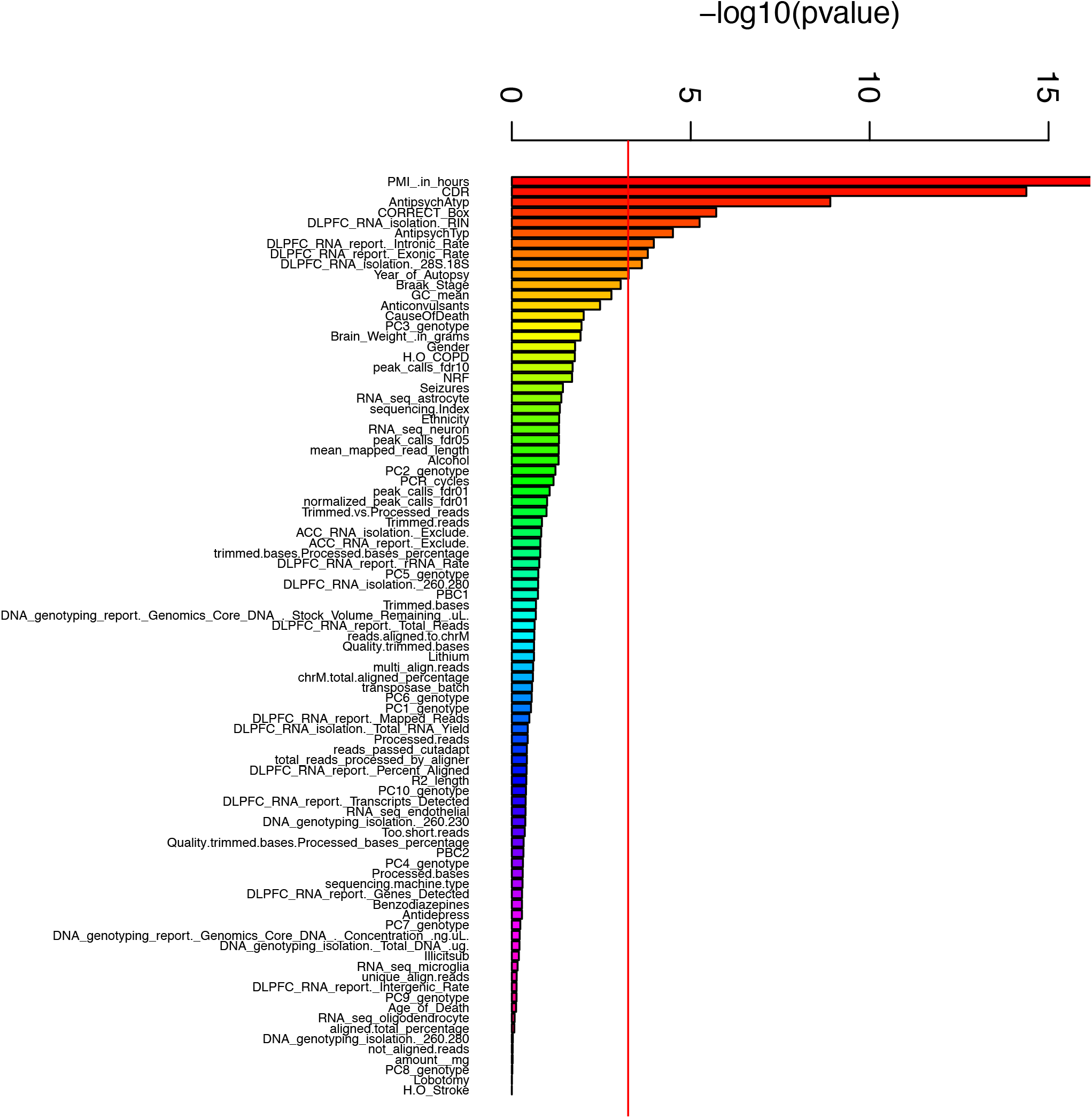
Association of metadata with case-control status. The associations between case-control status (N=135 cases, N=137 controls) and numerical variables were tested using linear regression, while χ tests were performed for categorical variables. The red bar represents the Bonferroni significance threshold (adjusted pvalue = 0.05).

**Figure S4:**
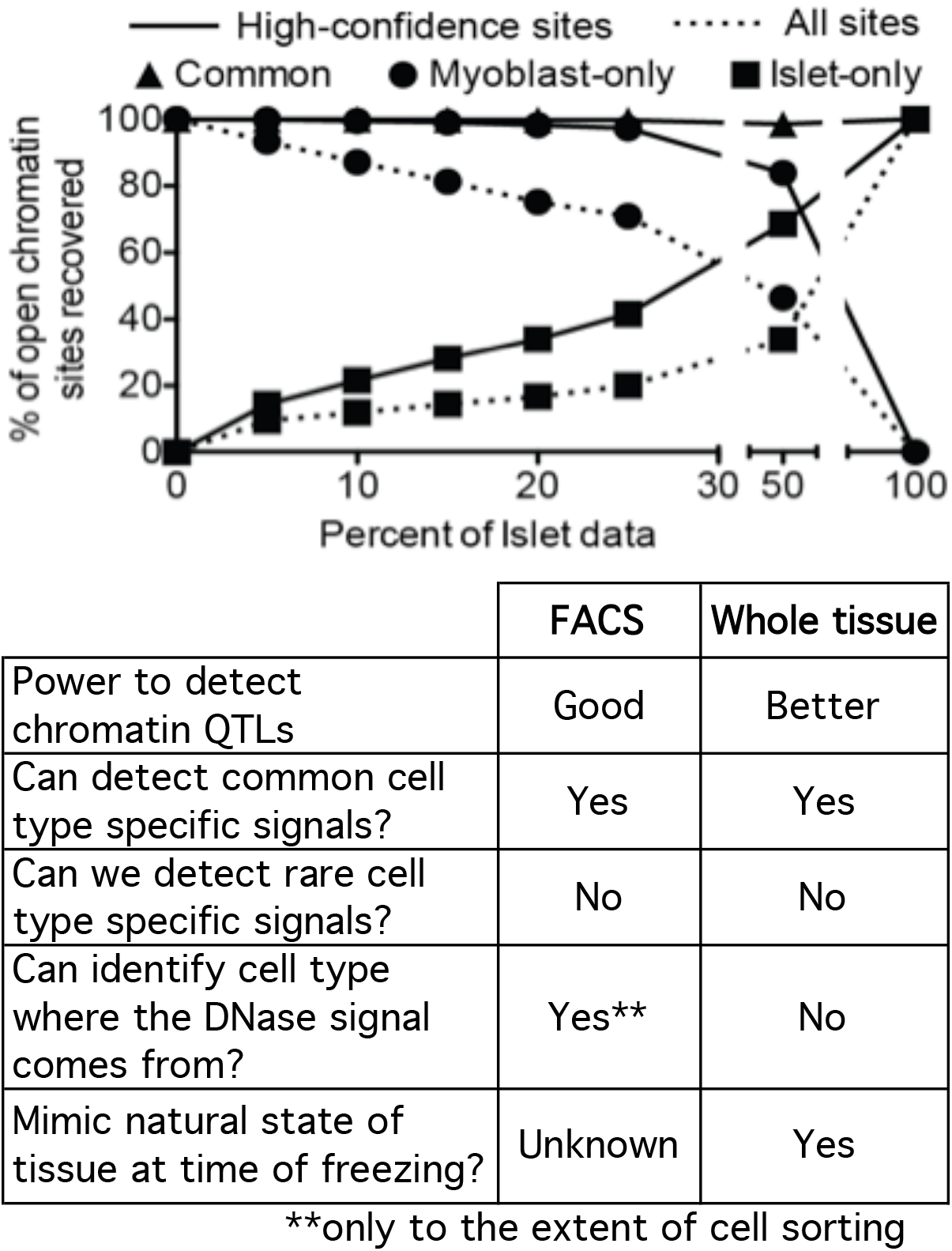
Identification of open chromatin in heterogeneous tissue. (A) DNase-seq data from myoblast and pancreatic islet were mixed in silico. The percentage of tissue-specific and shared chromatin regions are shown. (B) Advantages and disadvantages of FACS sorted nuclei vs. whole tissue analyses.

**Figure S5:**
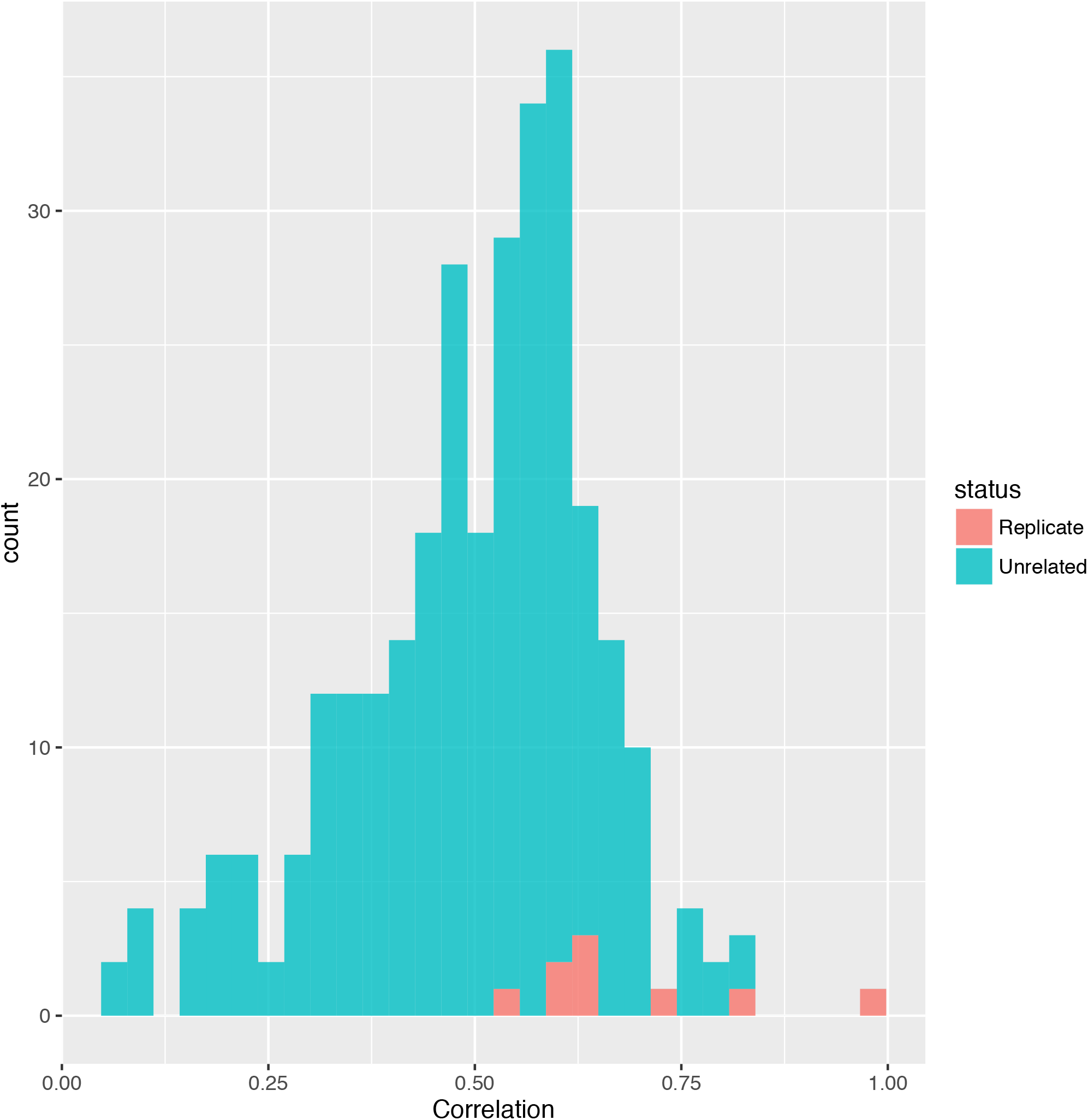
Spearman rank correlations of biological replicates. For biological replicates (9 pairs, red) and all possible combination of the sample with replicates with other samples excluding the replicate samples (blue). Pvalue=1×10^−4^.

**Figure S6:**
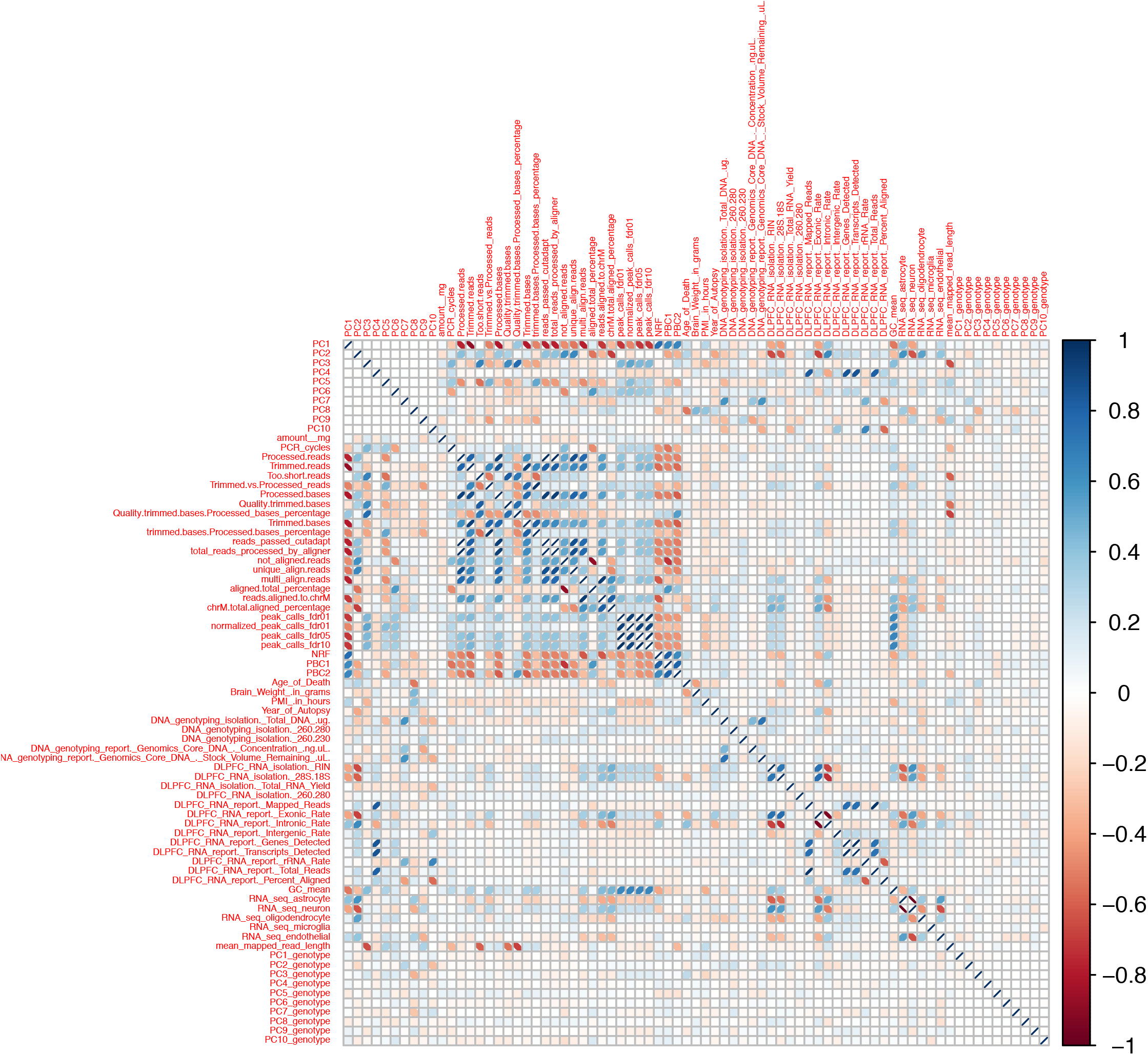
Correlation matrix for imputed numerical metadata in 288 samples. Pearson correlations were computed between all pairs of imputed numerical meta data variable (65 variables) with the addition of the 10 first principal components of the meta data variables. Blue shows positive correlations and red negative correlations.

**Figure S7:**
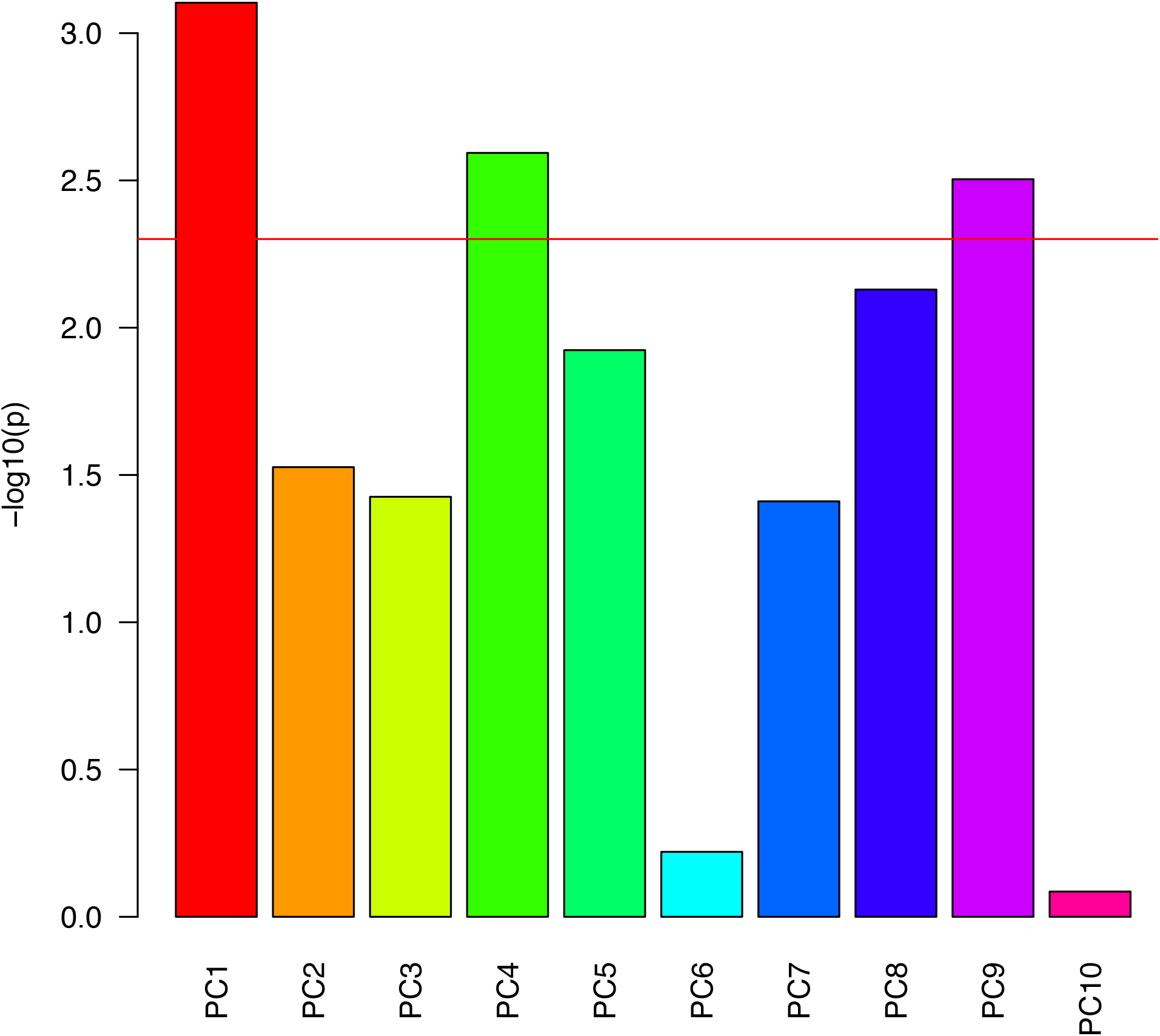
Association of PCs from imputed numerical metadata with diagnosis. The association pvalues were obtained by performing linear regressions. The red bar represents the Bonferonni significance threshold (adjusted pvalue = 0.05)

**Figure S8:**
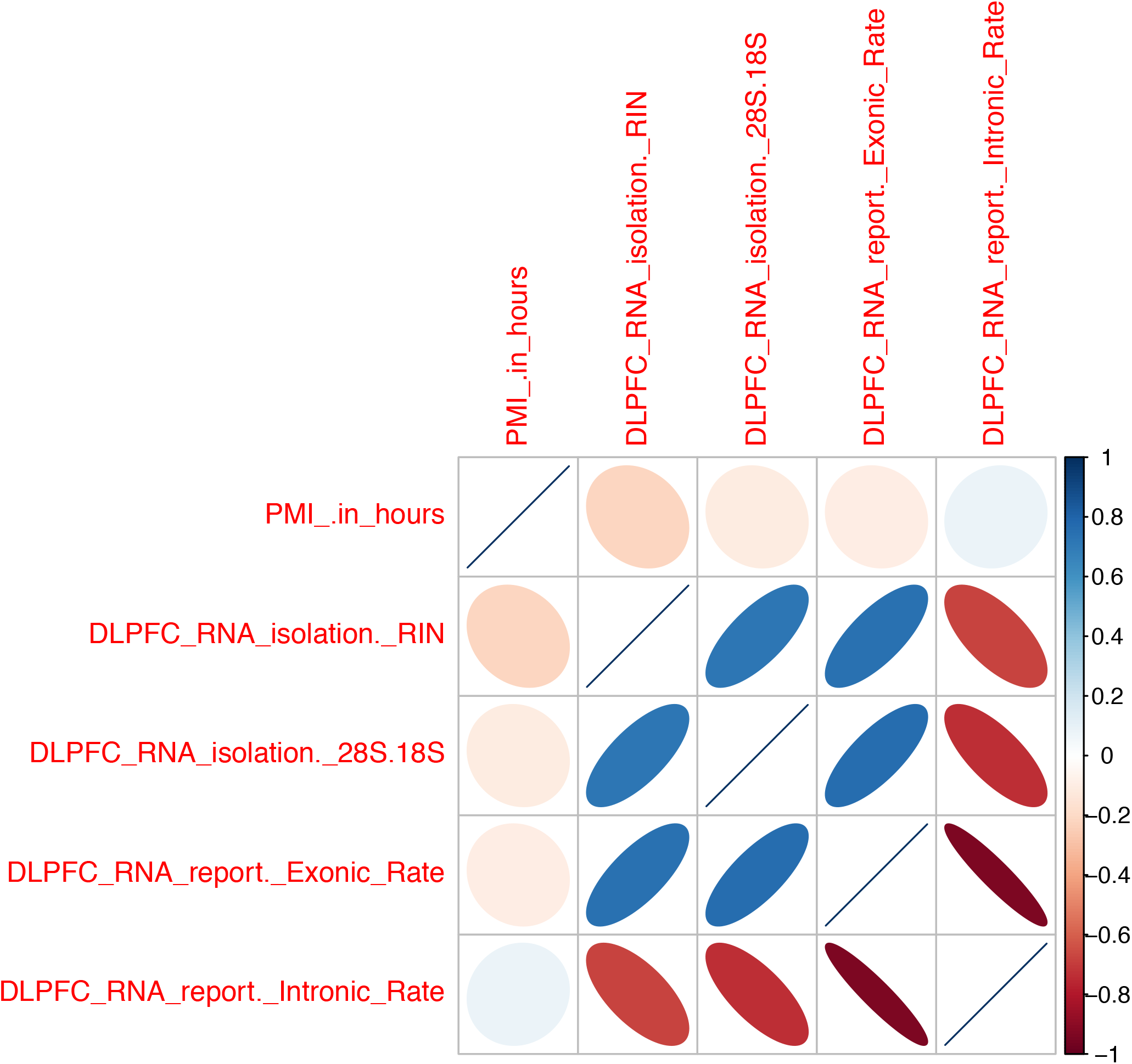
Intercorrelations of numerical metadata significantly associated with case-control status. Pearson correlation between all pairs of numerical meta data variables significantly associated with case/control status are represented (positive correlations in blue, negative correlations in red). Note: The RNA-seq meta data variables are strongly correlated together but are largely independent of the post-mortem interval.

**Figure S9:**
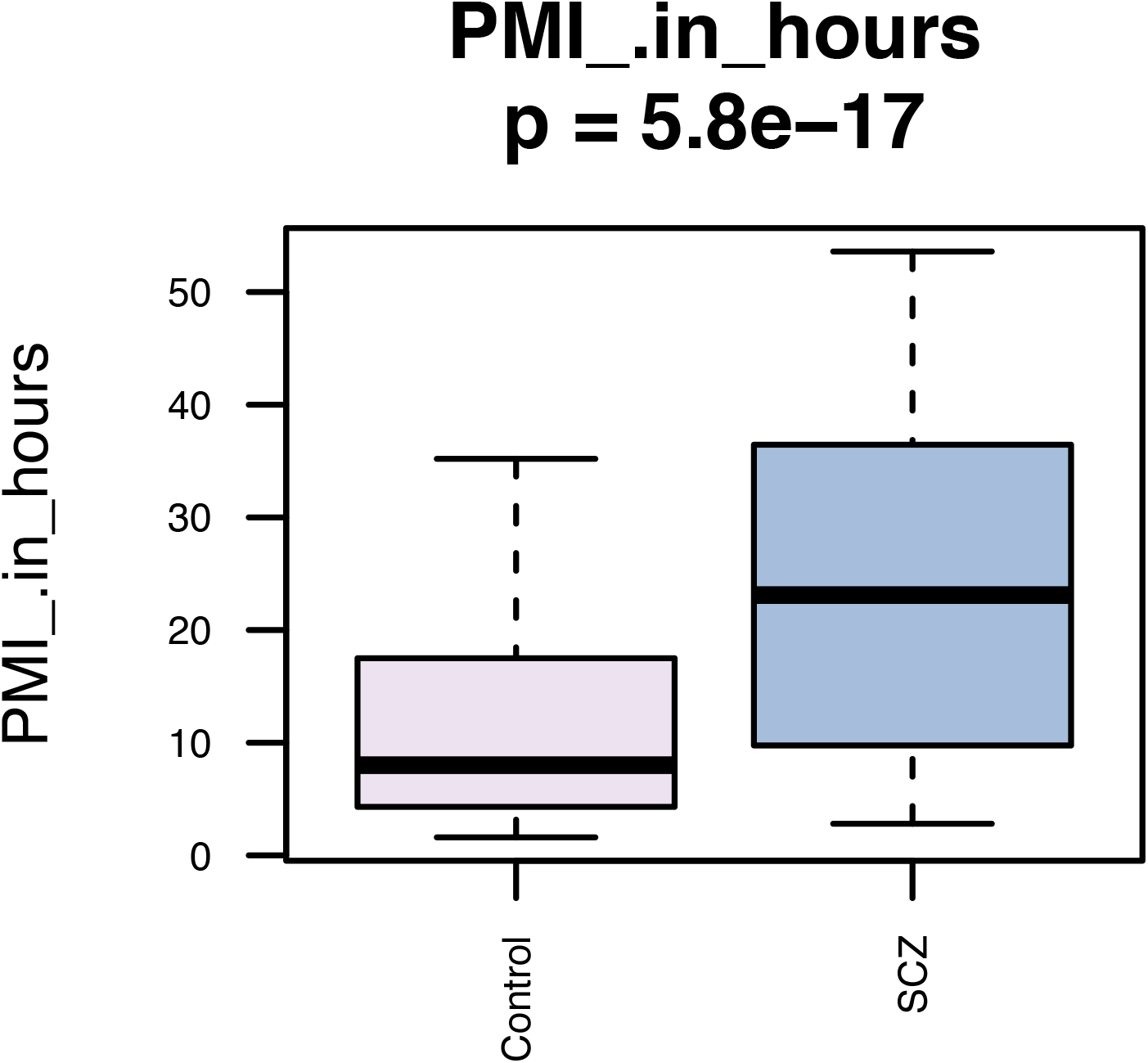
Postmortem interval and case-control status. The P-value was obtained by performing a linear regression between case-control status (N=135 cases, N=137 controls) and post-mortem interval.

**Figure S10:**
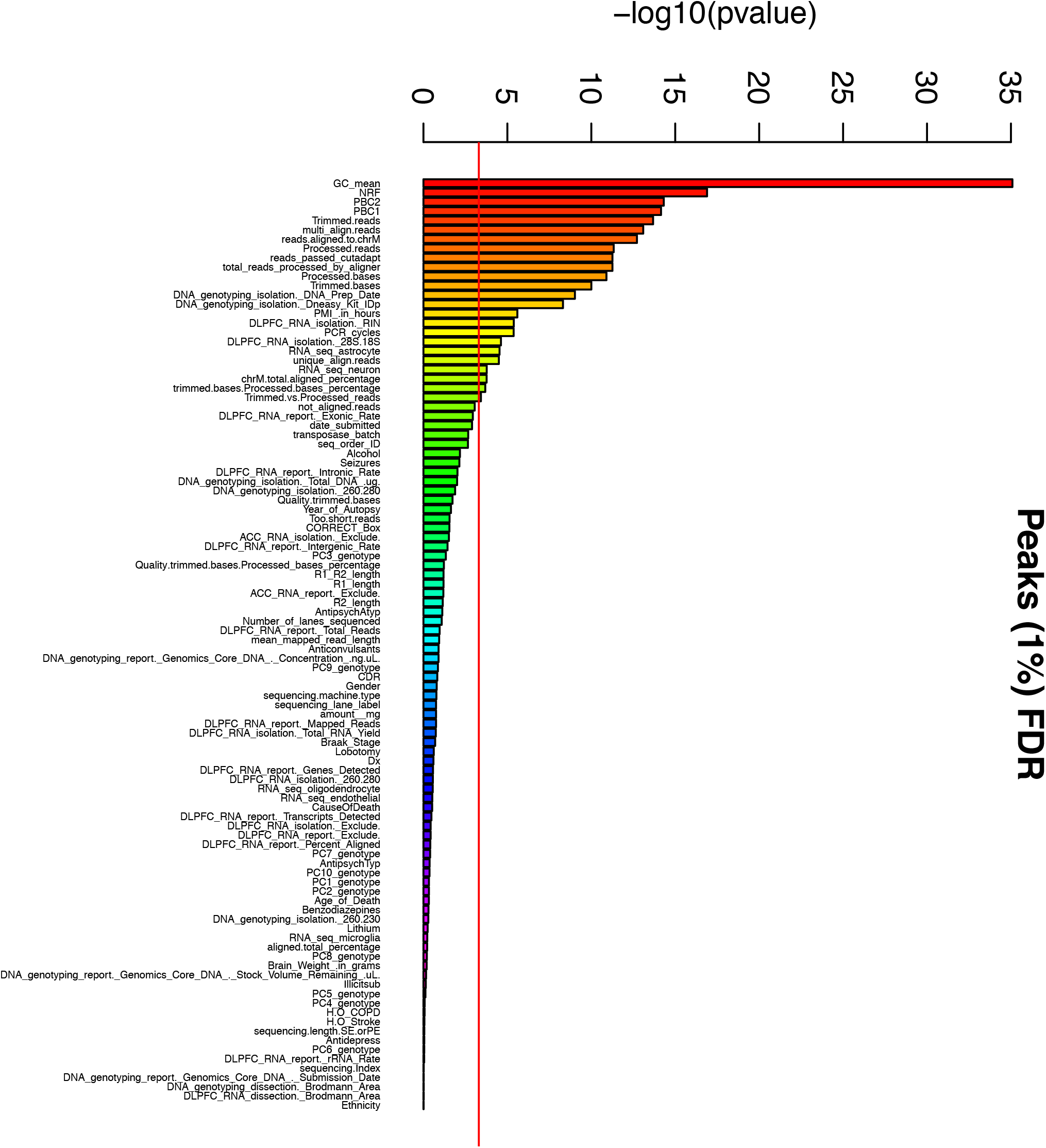
Association of number of ATAC-seq peaks (1% FDR) with metadata. The associations between the number of peaks (1% FDR) and numerical variables were tested using linear regression, while χ^2^ tests were performed for categorical variables. The red bar represents the Bonferroni significance threshold (adjusted pvalue = 0.05).

**Figure S11:**
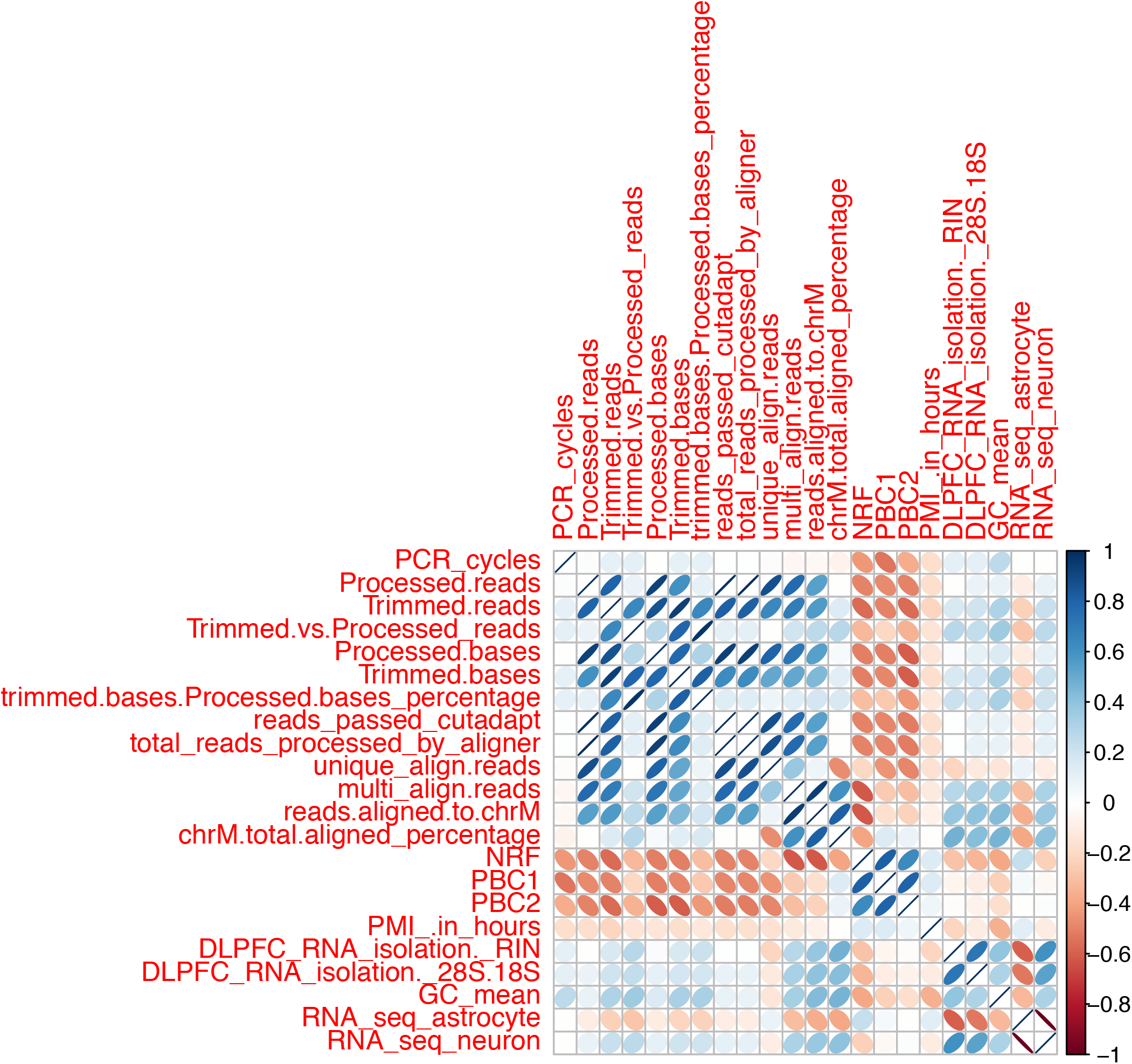
Correlation of technical covariates with number of peak calls. Pearson correlations of 22 numerical meta data variables significantly associated with the number of peak calls (1% FDR). Note: most of the meta data variables significantly associated with the number of peak calls are highly correlated.

**Figure S12:**
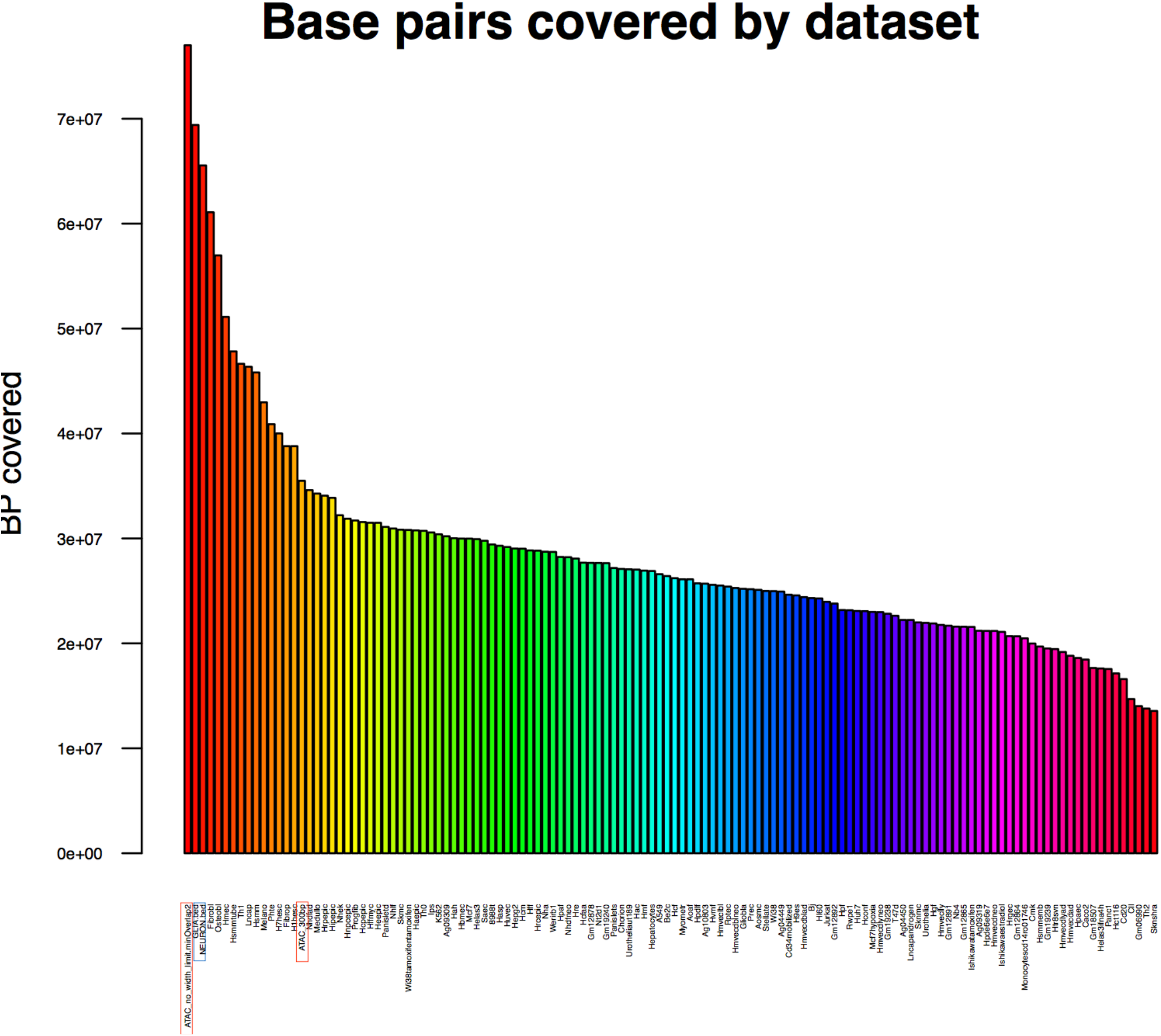
Coverage of ATAC-seq and DNase-seq datasets representing 125 cell types and tissues. The number of base pairs covered by significant peaks in each dataset is shown here. Note: without restricting our ATAC-seq peaks to 300bp, they would be covering significantly more base pairs then ENCODE DNase-seq datasets.

**Figure S13:**
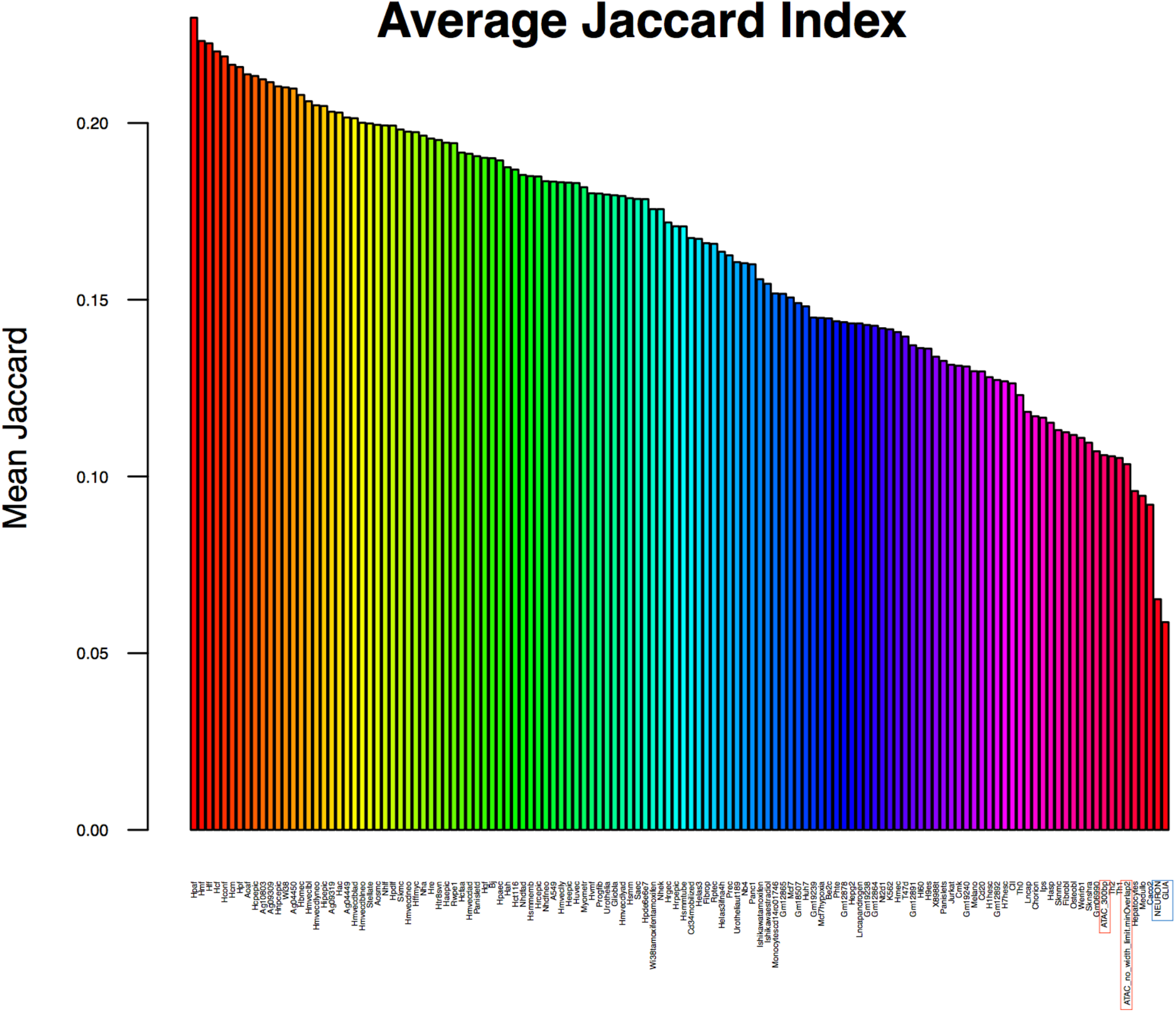
Mean Jaccard index for ATAC-seq and DNase-seq data in 125 cell types and tissues. The Jaccard index (intersection of two data sets divided by their union) was computed between all pairs of ATAC-seq or DNase-seq datasets. The average Jaccard index for each dataset is shown here.

**Figure S14:**
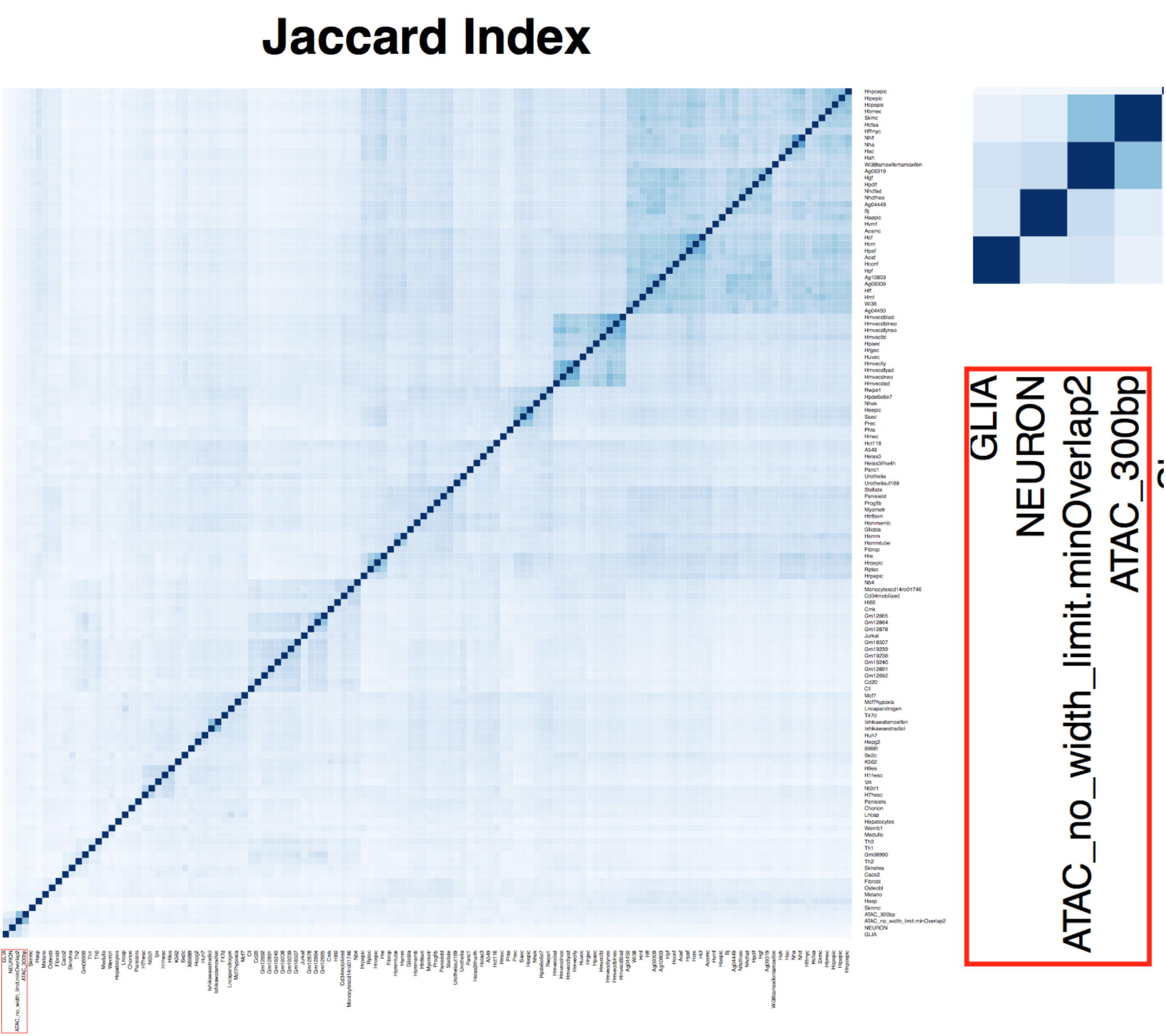
Jaccard index for ATAC-seq and DNase-seq data from 125 cell types and tissues. The Jaccard index (intersection of two data sets divided by their union) was computed between all pairs of ATAC-seq and DNase-seq datasets. White represents a Jaccard index of 0, while dark blue represents a Jaccard index of 1. Panel to right shows correlation similarity to ATAC-seq generated from sorted neuronal nuclei using NeuN+ antibody (NEURON) and NeuN-nuclei (GLIA).

